# Impaired Barrier Integrity of the Skeletal Muscle Vascular Endothelium Drives Progression of Cancer Cachexia

**DOI:** 10.1101/2022.12.12.520118

**Authors:** Young-Mee Kim, Mark A. Sanborn, Xinge Wang, Georgina Mancinelli, Sreeparna Chakraborty, Shaluah Vijeth, Priyanka Gajwani, Paul Grippo, Steve Seung-Young Lee, Tibor Valyi-Nagy, Peter T. Toth, Klara Valyi-Nagy, Jalees Rehman

## Abstract

Cancer patients experience cachexia, which is characterized by extensive skeletal muscle wasting that worsens the quality of life and increases mortality. Currently, there are no approved treatments that can effectively counteract cancer cachexia. Vascular endothelial cells (ECs) are essential for maintaining tissue perfusion, nutrient supply, and preventing inappropriate transmigration of immune cells into the tissue. However, little is known about the role of the muscle vasculature in cancer cachexia. We hypothesized that endothelial dysfunction in the skeletal muscle mediates cancer cachexia. Using transgenic pancreatic ductal adenocarcinoma (PDAC) mice and a tissue clearing and high-resolution 3D-tissue imaging approach, we found that the loss of skeletal muscle vascular density precedes the loss of muscle mass. Importantly, we show that cancer cachexia patients exhibit significantly decreased muscle vascular density and severe muscle atrophy when compared to non-cancer patients. Unbiased single cell transcriptomic analyses of the muscle endothelium unveiled a unique EC population present in cachexia muscles. Increased circulating Activin-A suppresses the expression of the transcriptional co-activator PGC1α in the muscle endothelium, thus disrupting junctional integrity in the vasculature and increasing vascular leakage. Conversely, restoration of endothelial-specific PGC1α prevented the decreased vascular density and muscle loss observed in tumor-bearing mice. Our study suggests that EC-PGC1α is essential for maintaining the integrity of the skeletal muscle vascular barrier and that restoring muscle endothelial function could be a valuable therapeutic approach to prevent or reverse cancer cachexia.

## INTRODUCTION

Cachexia is defined as skeletal muscle loss that can be observed in multiple chronic diseases including cancer, heart failure, chronic obstructive pulmonary disease, and chronic kidney disease^1^. The wasting of skeletal muscle significantly impairs the activities of daily living and is a key factor in determining the quality of life and mortality of patients^2^. Unlike starvation-induced sarcopenia, which can be reversed by subsequent supply of adequate nutrients and increased exercise^3^, cachexia is thought to result in predominantly irreversible muscle loss due to molecular and cellular dysfunction in the muscle tissue that persists even when nutrition is enhanced^2^. Cachexia is a multifactorial syndrome that has been attributed to a combination of factors present in chronic diseases such as inflammation, metabolic alterations, energy imbalance, and psychological factors^4–6^. As of now, there are no approved efficacious treatments for cachexia even though it is highly prevalent in advanced chronic diseases such as cancer^2, 4^. Recent research has focused on the impact of inflammatory cytokines such as IL-6 and TNF-α as likely mediators of cachexia^4, 7–9^. IL-6 has emerged as a central mediator of cachexia, which induces chronic inflammation^8^ and also directly impacts skeletal muscle^10^ but this raises intriguing mechanistic questions of how circulating inflammatory mediators or circulating immune cells enter the cachexic muscle to promote cachexia and how additional cell populations or mediators contribute to cachexia.

Endothelial cells (ECs) cover the inner walls of blood vessels and thereby are directly exposed to circulating immune cells, nutrients, growth factors, cytokines and other small molecules in the blood^11, 12^. Therefore, ECs are highly responsive and sensitive to changes in properties of the blood and regulate organ and tissue homeostasis with functional heterogeneity dependent on states, vascular beds, and tissues^13–16^. Skeletal muscle accounts for up to 40% of total body weight in healthy individuals and is composed of several different types of cells, including mature myocytes which form muscle fibers, regenerative muscle satellite cells, and ECs^17^. Importantly, skeletal muscle is a highly vascularized tissue^18^ due to the continuous need for oxygen and nutrients supplied by the blood via the perfused microvascular network. By maintaining an intact vascular barrier, ECs prevent the uncontrolled leakage of circulating molecules or inappropriate transmigration of circulating immune cells into the interstitium^19, 20^. Although the role of the muscle vasculature in mediating cachexia has not yet been studied in depth, there are some important clues derived from aging that point to a potential mechanistic role for ECs in cachexia progression. Aging in humans is associated with a loss of muscle vasculature and this loss of vasculature is correlated with a decrease in muscle mass and muscle strength in the elderly^21, 22^, mirroring key features of cancer cachexia^23^ and thus suggesting that similar mechanisms could be at play. Importantly, the affected muscles are often spatially removed from active disease sites (such as in the setting of abdominal tumors or cardiomyopathy) indicating that cachexic factors are likely released from disease sites and can remotely act on the muscle via the blood circulation. This highlights the importance of ECs which are the first point of contact for circulating factors in the blood. However, it is not known how these circulating factors may affect remote ECs.

The transcriptional co-activator PGC1α (peroxisome proliferation activator receptor-γ coactivator1-α) is a transcriptional coactivator which acts as a key regulator of oxidative metabolism and redox homeostasis in various tissues such as heart, liver, brown adipose tissue, skeletal muscles, and brain^24, 25^. PGC1α expressed in skeletal myocytes is required for muscle endurance^26, 27^. Overexpression of PGC1α in myocytes can partially suppress age-related sarcopenia^25^ and in cancer cachexia^28^. More recently, PGC1α has emerged as an important regulator of endothelial function. Expression of endothelial PGC1α reduces vascular dysfunction^29, 30^ and mediates compensatory or adaptive angiogenesis in disease conditions where blood supply is compromised^31, 32^.

In this study, we investigated molecular mechanisms of cancer cachexia using two complementary *in vivo* tumor models: 1) KPC (*LSL-KrasG12D/+;LSL-Trp53R172H/R172H;Pdx-1-Cre)* mice^33^ which develop malignant pancreatic tumors and phenocopy several features of human pancreatic ductal adenocarcinoma (PDAC), a leading cause of cancer cachexia^2^ and 2) a mouse melanoma allograft model which we found to exhibit a rapid concomitant tumor and cachexia progression. Using both cancer cachexia mouse models, we investigated the temporal and mechanistic relationship between skeletal endothelial dysfunction and the muscle loss in cancer cachexia as well as the potential whether changes of muscle vascular density precede muscle loss in cancer cachexia and whether circulating cachexic cues induce muscle vascular dysfunction. Furthermore, we assessed the benefits of therapeutically overexpressing PGC1α in the muscle endothelium of tumor bearing mice.

## RESULTS

### Microvascular density is decreased in cancer cachexic muscles

To define the role of muscle vascular endothelial function in cancer cachexia, we used transgenic KPC (LSL-**K**ras^G12D/+^: LSL-Tr**p**53^R172H/R172H^: Pdx1-**C**re) mice which were generated by crossing LSL-Kras^G12D/+^: LSL-Trp53^R172H/ R172H^ mice with Pdx-1 (pancreas duodenum homeobox transcription factor)-Cre mice^33, 34^. KPC mice spontaneously develop fibrotic pancreatic tumors and cachexia at approximately 4-6 months of age. Therefore, we evaluated the KPC-PDAC mice from 3 months onward. The KPC-5 month mice had >10% less body weight than age-matched control wild-type mice (WT-5mon), whereas no such difference was seen at 3 months (**Extended Data Fig. 1a**). Thus, we referred to these KPC mice which developed cachexia at 5 months as KPC-cachexia mice. First, we evaluated muscle endothelial density in intact muscle tissues using transparent tissue tomography (T3) method^35, 36^. The T3 method uses aqueous sugar solution-based immersion tissue clearing to enable high-resolution three-dimensional (3D) microscopy of the microvasculature in muscle tissues. The gastrocnemius muscles from control (WT, 5 month) and KPC-cachexia (5 month) mice were harvested, and immediately fixed and processed for tissue clearing with a fructose gradient followed by immunofluorescence staining using the endothelial marker CD31. The muscle capillary networks were intact and continuously connected in control mice, whereas they were significantly diminished and fragmented in KPC-cachexia mice (**Fig. 1a**). In addition, KPC-cachexic muscles exhibited significantly decreased the surface volume of vasculature compared to control muscles (**Fig. 1b**). We next quantified the number of ECs in the tibialis anterior (TA), gastrocnemius (GC), and quadriceps, which are major muscles important for muscle endurance^37, 38^. ECs from mixed skeletal muscles of hind limbs were isolated by enzymatic digestion and stained with an antibody against the EC marker CD31 as well as co-stained for the leukocyte marker CD45 to exclude small myeloid cell subsets which co-express CD31 and could inflate EC numbers. We also stained with DAPI to exclude dead cells and performed flow cytometry analysis. Consistently, the muscles of KPC-cachexic mice had a significantly decreased number of muscle ECs compared to control mice (**Fig. 1c**).

**Figure 1.**
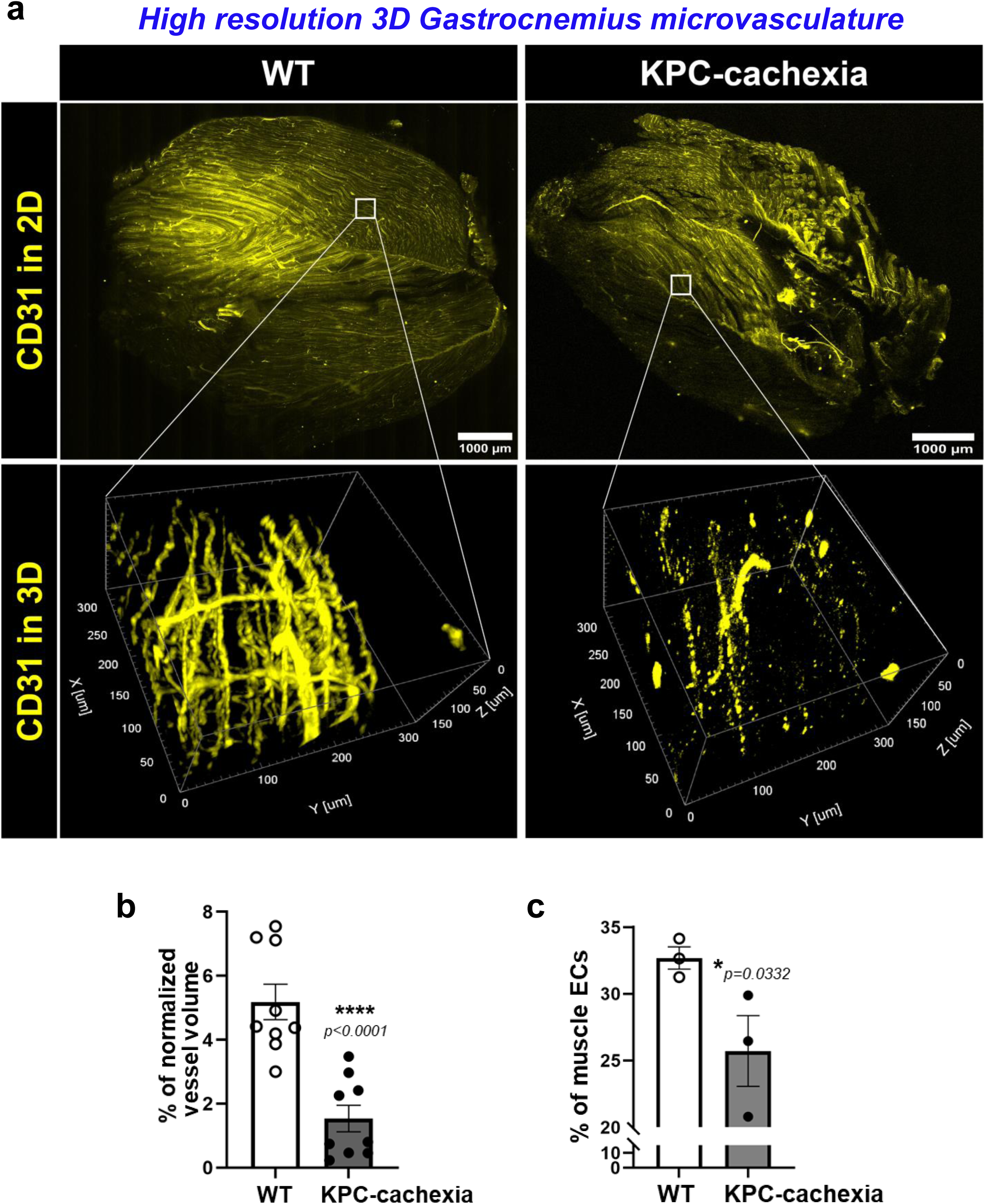
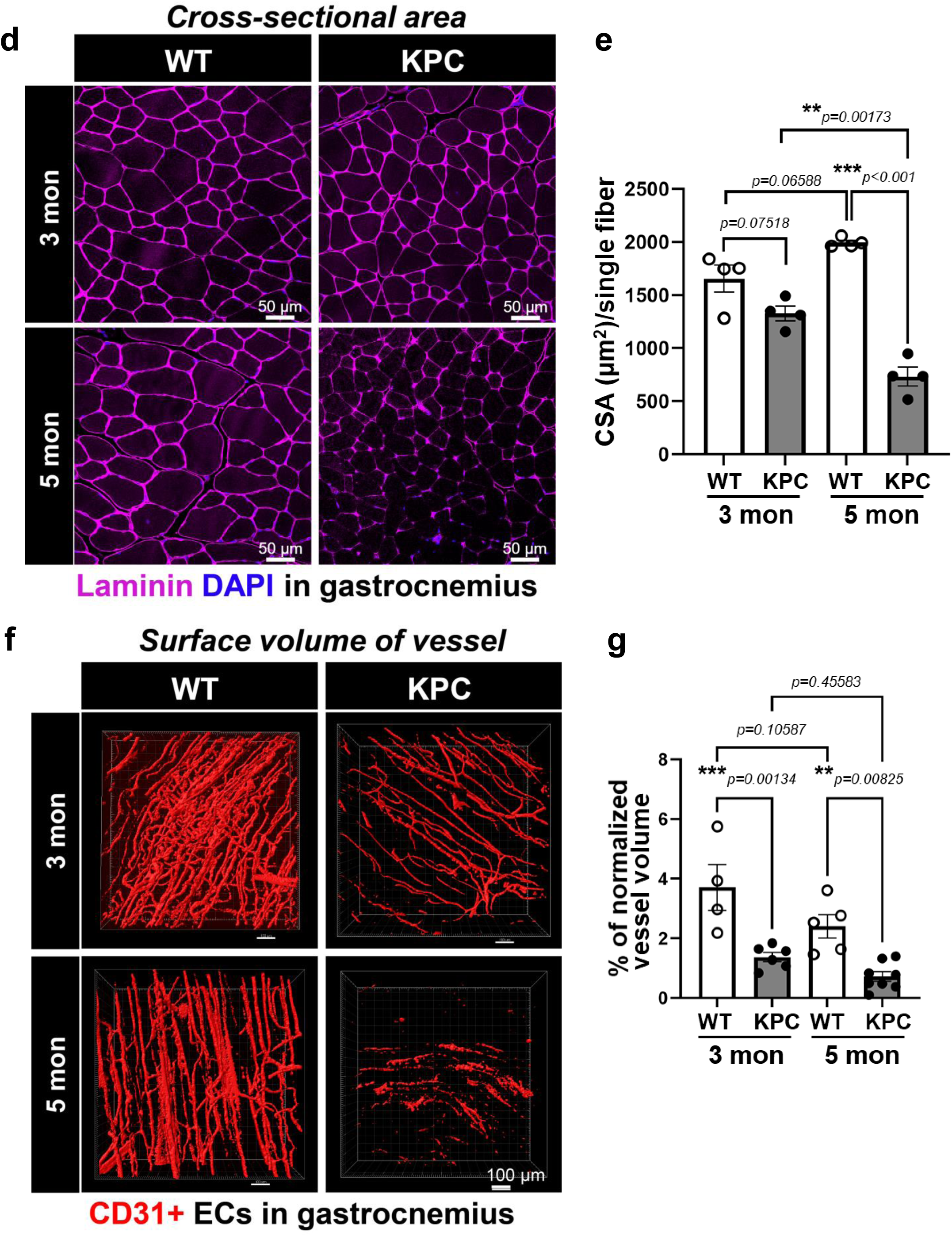
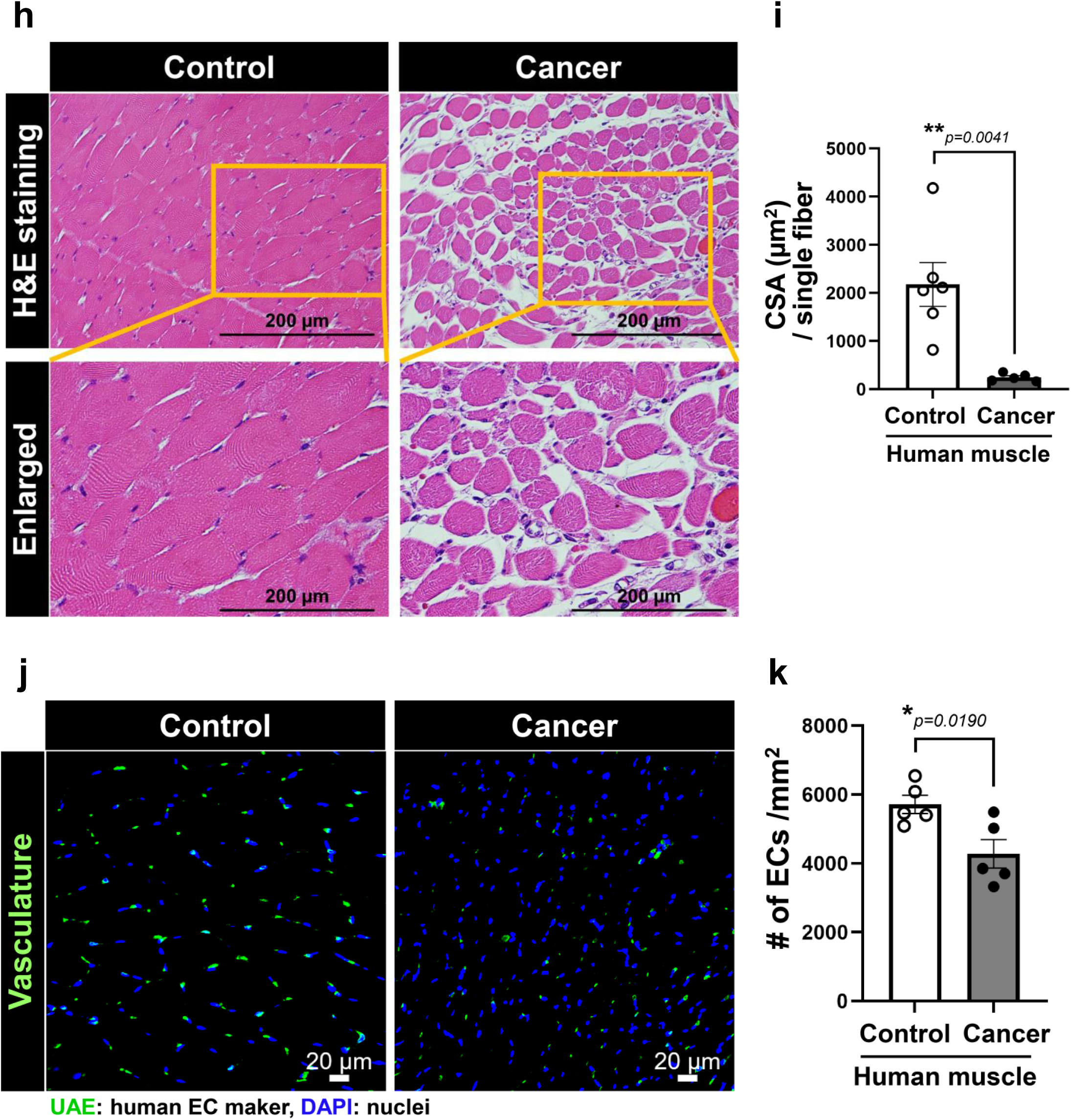
Muscle vascular density is decreased in cancer cachexic muscles. **a.** The gastrocnemius (GC) muscles from WT and KPC-cachexia (5 month) were briefly fixed, longitudinally sectioned with 400 µm, stained for fluorescent anti-CD31 (yellow) antibody, and optically cleared in fructose solutions. The 3D images were taken using a confocal fluorescence microscope (Caliber ID, RS-G4 with a 40x oil objective) and reconstructed for visualization of muscle microvasculature using Imaris software. 2D vasculature images (upper panel) in whole muscle tissue and 3D vasculature images (lower panel) in selected areas (white square box) were taken with x20 and x40 objectives, respectively. High resolution 3D images were obtained from 9 different area images in a muscle tissue (n=6 for each group). The representative 3D image data were shown. **b.** Muscle vasculature density was presented with percent (%) of normalized CD31+ vessel volume (µm^3^) in total muscle volume (µm^3^). Data are mean±SE from n=6 mice. The *p* values were evaluated by unpaired, two-tailed t-test. **c.** Quantification of ECs in mixed skeletal muscles (TA, GC, and quadriceps) by FACS analysis. The CD31**+**CD45**-**DAPI**-** cells were considered to be live pure ECs and presented with percent (%) of ECs in total muscle cells. Data are mean±SE for n=3 mice. The *p* values were evaluated by paired, two-tailed t-test. **d.** Cross-sectioned gastrocnemius muscle from WT mice (3 and 5 month) and KPC mice (3 and 5 month) were stained with laminin antibody for muscle base membrane (magenta). The images were taken using confocal microscopy (LSM880, 20x objective). The nuclei were visualized with DAPI. **e.** Muscle size in **d** was determined with cross sectional area (CSA) per single fiber in laminin-stained muscle using ImageJ. Data are mean±SE from n=4 mice. The statistical analysis was performed by one-way ANOVA, followed by post-hoc multiple comparison analysis using Tukey test, and the adjusted p-values were reported. **f.** The gastrocnemius (GC) muscles of WT mice (3 and 5 month) and KPC mice (3 and 5 month) were briefly fixed, tissue cleared, longitudinally sectioned with 400 µm and stained for CD31(red) specific antibody. The images were taken using high resolution 3D confocal microscopy with 40x objectives (Caliber ID RS-G4) and reconstructed for visualization using Imaris software. The surface volume of CD31+ area in 3D vasculature images was calculated based on the surface reconstruction of CD31+ signal using Imaris x64 7.7.2 software. The images are representative of n=3 mice. The scale bar is 100 µm. **g.** Muscle vasculature density was presented with percent (%) of normalized CD31+ vessel volume (µm^3^) in total muscle volume (µm^3^). Data are mean±SE from 4-8 different fields from n=3 mice. The statistical analysis was performed by one-way ANOVA, followed by post-hoc multiple comparison analysis using Tukey test, and the adjusted p-values were reported. **h-k.** Abdominal muscles from control patients and cancer cachexia patients were used to evaluate muscle size and vasculature. General information of patients can be found in the Supplementary information. **h.** Structure of muscles was showed with H&E staining. **i.** The cross-sectional area per single fiber was measured with Image J. Data are mean±SE from control patients (n=6) and cancer cachexia patients (n=5), and average n=30 single fiber from each muscle section. The *p* values were evaluated by unpaired, two-tailed t-test. **j.** Muscle endothelial cells were visualized by staining with human EC specific UEA1 and DAPI for nucleus. The images were taken using confocal microscopy (LSM880, 20x objective). **k.** Muscle vascular density was determined by counting number of UAE+ cells per muscle area (mm^2^). Data are mean±SE from control patients (n=5) and cancer cachexia patients (n=5), and 5-6 different fields from each muscle section. The *p* values were evaluated by unpaired, two-tailed t-test.

To determine whether muscle mass was correlated with muscle vascular density in cancer cachexia progression, we compared muscle size and muscle vascular density in gastrocnemius muscles from non-cachexia KPC-3 mon and cachexia KPC-5 mon mice as well as age matched WT mice. The muscle basal membrane and muscle ECs were stained with antibodies targeting laminin and CD31, respectively. Muscle size was determined with cross sectional area (CSA) per single fiber. The muscle cross sectional area decreased in KPC-cachexia (5mon) when compared to age-matched controls but such a decrease was not yet present in 3 month old KPC mice (**Figs. 1d-e**). Moreover, KPC-cachexia mice had significantly higher numbers of small fibers (<500 µm^2^) as well as fewer large fibers (1500-2500 µm^2^) when compared to control mice (**Extended Data Fig. 1b**). Interestingly, the vascular density was already decreased at the age of 3 months in the KPC mice and showed only limited further decrease, suggesting that the vascular rarefaction preceded the loss of muscle mass (**Figs. 1f-g**). Importantly, increased expression of *MuRF1* and *Atrogin1*, E3 ubiquitin ligases which are typically upregulated during muscle atrophy and serve as markers of cachexia^39^ was seen only in KPC-5mon but not KPC-3mon mice, consistent with the notion that cachexic muscle loss occurs at a more advanced age than the early loss of vessels in the muscle (**Extended Data Fig. 1c**).

We examined whether loss of muscular vasculature in cancer cachexia affects functional performance by measuring mouse grip strength and fatigue which are indicative of whole-body muscle strength in mice^40^. We found a significant decrease in mouse grip strength in KPC-5mon, but not KPC-3mon mice (**Extended Data Fig. 1d**). Similarly, increased fatigue was also observed in KPC-5mon, but not KPC-3mon mice (**Extended Data Fig. 1e**). These results support the notion that decrease of muscle vascular density in KPC mice occurs independently of aging and precedes the loss of muscle mass and function.

To determine the translational relevance of muscle vascular rarefaction in cancer, we evaluated autopsied abdominal muscles from non-cancer patients (referred to as controls) and cancer patients who had developed cachexia. The muscles of patients with a diagnosis of cancer cachexia exhibited very severe muscle atrophy as evidenced by the smaller and more irregular muscle fiber CSA compared to control patients (**Figs. 1h-i** and **Extended Data Fig. 1f**). Moreover, cancer cachexia patients also had markedly decreased muscle vascular density as shown by the human EC marker, Ulex Europaeus Agglutinin-I (UEA1) (**Figs. 1j-k**). These data suggest that loss of muscle vasculature also occurs in cancer patients.

### Vascular endothelial apoptosis and endothelial to mesenchymal transition are increased in skeletal muscle of tumor bearing mice prior to the onset of muscle loss

Next, we explored the mechanisms governing the decreased muscle vascular density and discontinuous or fragmented vascular networks seen in KPC cachexia mice. To address whether the loss of muscle vasculature was due to cell death, sections of the gastrocnemius muscles of WT and KPC-mice were assessed using an *in situ* TUNEL (**T**erminal deoxynucleotidyl transferase d**U**TP **n**ick **e**nd **l**abeling) assay. Interestingly, KPC-3mon mice demonstrated more endothelial apoptosis than WT (5mon) mice as shown in **Figs. 2a-b**, and the number of apoptotic ECs was indeed increased in the muscle vasculature from KPC-3mon mice which do not yet show overt evidence of cachexia. We hypothesized that circulating factors released by the tumor could affect the remote muscle vasculature because muscle capillary vessels respond to factors in the blood. Activin, a member of the transforming growth factor beta (TGFβ) superfamily which regulates cell differentiation, is elevated in the blood in multiple chronic diseases^41–44^. Activin-A has been reported to suppress EC proliferation under hypoxia^45^ suggesting the possibility that Activin-A acts as a negative regulator of muscle vasculature. We therefore measured Activin-A levels in the blood plasma of control and KPC-cachexia mice. KPC-cachexia mice exhibited higher levels of circulating Activin-A than control mice (**Fig. 2c**). Ex vivo exposure of human microvascular ECs (HMVECs) to Activin-A (25 ng/mL) significantly increased apoptosis (**Fig. 2d**) and was accompanied by decreased expression of *Mcl1*, an anti-apoptotic gene of the Bcl-2 family using RT-qPCR (**Fig. 2e**). In addition, Activin-A treated ECs had significantly increased mRNA levels of endothelial-to-mesenchymal transition (EndMT)^16^ markers *Snail* and *Vimentin* compared to vehicle treated ECs (**Fig. 2f**). Thus, we examined whether EndMT in muscle ECs was induced during cachexia progression in KPC mice by co-staining with EC marker isolectin B4 (IB4) and mesenchymal marker α-smooth muscle actin (SMA). We observed a significant increase in α-SMA area was observed in muscle ECs of KPC-3mon mice compared to control mice (**Figs. 2g-h**). The isolated muscle ECs from KPC-cachexia mice had significantly increased mRNA levels of EndMT marker *Twist* relative to WT (**Extended Data Fig. 2a**). Taken together, our results suggest that EC apoptosis and EndMT in muscle vasculature contribute to the loss of functional muscle vasculature and precede the loss of muscle mass or muscle function.

**Figure 2.**
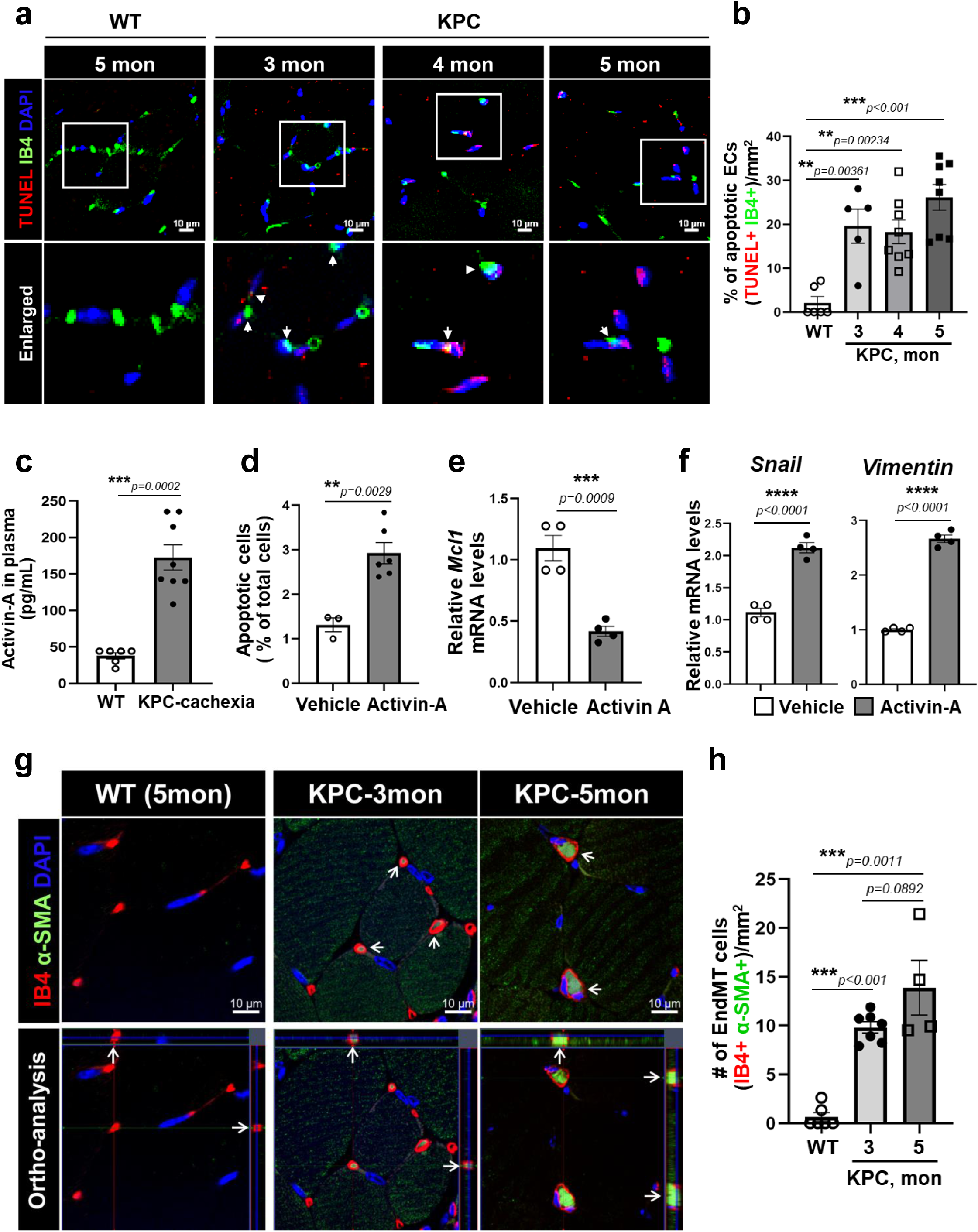
Decrease of muscle vascular density was caused by endothelial death and differentiation from endothelial to mesenchymal cells in cancer cachexia progression. **a.** Cross-sectioned gastrocnemius muscle from WT mice (5 month) and KPC mice (3, 4, and 5 month) were used *in situ* TUNEL assay for apoptotic cells (red) and co-stained with IB4 for ECs (green). The images were taken using confocal microscopy (LSM880, 20x objective). The nuclei were visualized with DAPI. **b.** Number of double positive cells of TUNEL (red) and IB4 (green) in total ECs (IB4+) in **a** was quantified by Image J and the apoptotic ECs were presented with percent (%) in total ECs per field (mm^2^). Data are mean±SE from 5-13 different fields from n=3 mice for each group. The statistical analysis was performed by one-way ANOVA, followed by post-hoc multiple comparison analysis using Tukey test, and the adjusted p-values were reported. **c.** The blood plasma from WT mice and KPC-cachexia mice (5 months) was used to measure Activin-A levels using ELISA Kit. Data are mean ± SE for n=7-10 mice. The *p* values were evaluated by unpaired, two-tailed t-test. **d-f.** The HMVECs were treated with Activin-A (25 ng/mL) for 24 h. **d.** The cell apoptosis was determined by staining with Annexin-V-FITC and PI following FACS analysis. The apoptotic cells (Annexin V-FITC+PI+) were presented with percent (%) of total cells. Data are mean±SE for n=3-6 biological replicates. The *p* values were evaluated by unpaired, two-tailed t-test. **e.** The expression of anti-apoptotic gene, *Mcl1* was determined by RT-qPCR with specific primers. The levels of *Mcl1* were normalized by *18s* levels and presented fold change. Data are mean±SE for n=4 biological replicates. The *p* values were evaluated by unpaired, two-tailed t-test. **f.** The expression of endothelial to mesenchymal transition (EndMT) genes, *Snail* and *Vimentin* was determined by RT-qPCR with their specific primers. The levels of *Snail* and *Vimentin* were normalized by *18s* levels and presented fold change. Data are mean±SE for n=4 biological replicates. The *p* values were evaluated by unpaired, two-tailed t-test. **g-h.** Cross-sectioned gastrocnemius muscle from WT mice (5 month) and KPC mice (3 and 5 month) were stained with IB4 for ECs (red) and with α-SMA antibody for smooth muscle actin (green). The images were taken using confocal microscopy (LSM880, 63x objective). The nuclei were visualized with DAPI. **h.** Quantification of EndMT cells in **g** was presented with number of double positive cells of IB4 (red) and α-SMA (green) in total ECs (IB4+) per muscle area(mm^2^). Data are mean±SE from 4-7 different fields from n=3 mice. The statistical analysis was performed by one-way ANOVA, followed by post-hoc multiple comparison analysis using Tukey test, and the adjusted p-values were reported.

Next, we investigated whether the observed loss of muscle vascular density prior to the onset of cachexia was specific to the KPC model of pancreatic adenocarcinoma, or also present in other forms of cancer cachexia by studying a melanoma allograft cachexia model. Mouse skin melanoma B16F10 cells were implanted subcutaneously in the dorsal flank of C57/BL6 mice. We harvested major skeletal muscles from the hindlimbs in a time dependent manner (**Extended Data Fig. 2b**) and evaluated muscle vascular density compared to control mice. The muscle vascular density in cross-cryosections was significantly decreased in melanoma bearing mice compared to control mice, as visualized by IB4 staining for ECs (**Extended Data Figs. 2c-d**) and the number of muscle ECs of melanoma bearing mice also significantly decreased from week 1 onwards as compared to control mice (**Extended Data Fig. 2e**). We did not find any tumor metastases in the examined skeletal muscle, suggesting that the muscle wasting was not due to tumor metastasis into the local muscle, but likely due to remote effects of the tumor. Taken together, our data indicate decreased muscle vasculature density precedes muscle loss in major muscles of two experimental models of cancer cachexia.

### Cachexic muscles exhibit leaky capillaries and an increase in immune cell infiltration

We next assessed endothelial barrier integrity as a key read-out of intact vascular function in tumor bearing mice. Control and melanoma bearing mice (3 weeks) were injected intravenously with FITC-albumin dye. After circulating FITC-albumin was removed by PBS perfusion, the gastrocnemius muscles were immediately harvested. Cryosections of 50 µm thickness were evaluated for the presence of FITC-albumin fluorescence by confocal microscopy (**Fig. 3a**). The vasculature of the gastrocnemius muscles in melanoma bearing mice had significantly higher microvascular leak than the muscles of control mice (**Fig. 3b**), indicating that the presence of a tumor at remote sites can increase muscle vascular leakage. To examine whether such leakiness would compromise oxygen transport into the tissue – a central function of the muscle vasculature-we injected the mice intraperitoneally with FITC-pimonidazole which specifically binds to the thiol group of proteins or peptides in hypoxic cells or regions and thus serves as a marker of tissue hypoxia. The muscles of melanoma bearing mice showed significantly increased hypoxia (**Figs. 3c-d**), indicative of reduced perfusion by the rarified and leaky muscle vasculature. The tibialis anterior muscles of melanoma bearing mice also had significantly increased expression of the glucose transporter *Glut1* which is increased as a compensatory adaption to hypoxia to facilitate anaerobic glycolysis (**Fig. 3e**).

**Figure 3.**
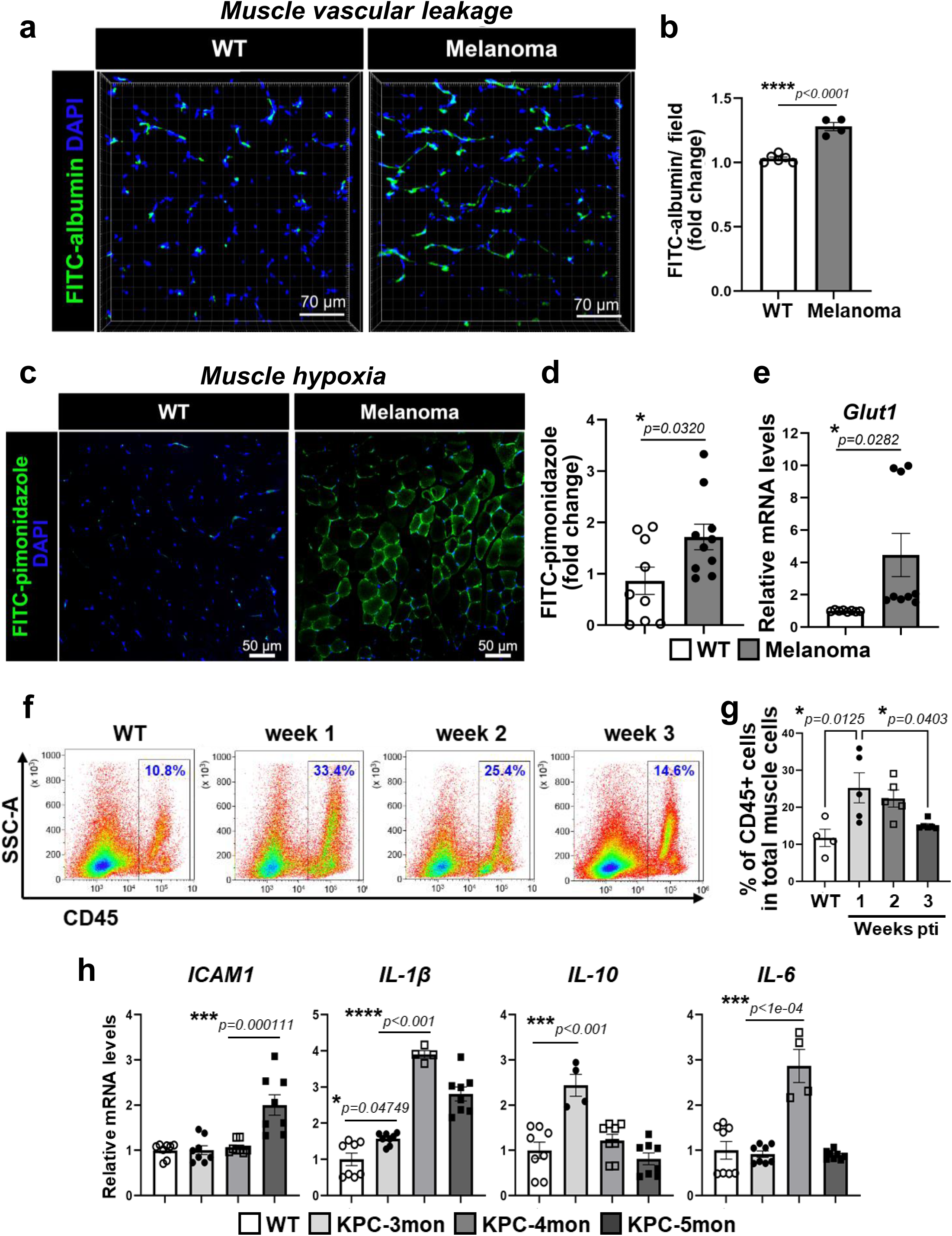
Decreased muscle vascular density is associated with increased vascular leakiness and immune cell infiltration into muscles of tumor bearing mice. **a-b.** WT and melanoma bearing mice for 3 weeks were retro-orbitally administrated with FITC-albumin (250 mg/kg in PBS). After 10 min, circulating FITC-albumin was washed by PBS perfusion. **a.** The FITC-albumin fluorescence images in gastrocnemius muscle cryosections (50 µm) were taken by Z-sectioning using confocal microscopy (LSM710, 20x objective) and reconstructed for visualization using Imaris software. Nucleus was stained with DAPI. The representative images are from n=4-6 mice. **b.** Quantification of fluorescence intensity of FITC-albumin in **a** using image J. Data are mean±SE for n=4-6 mice. The *p* values were evaluated by unpaired, two-tailed t-test. **c-e.** WT and melanoma bearing mice for 3 weeks were intraperitoneally received with hypoxic probe pimonidazole (100 mg/kg) 1h before harvesting muscle and the extent of hypoxia in gastrocnemius muscle was determined by the Hypoxyprobe Plus Kit. Nucleus was stained with DAPI and the images were taken using confocal microscopy (LSM880, 20x objective). **c.** The representative images are from n=4-5 mice. **d.** The FITC-pimonidazole fluorescence intensity in **c** was measured using image J. Data are mean±SE for n=9-10 different fields from n=4-5 mice. The *p* values were evaluated by unpaired, two-tailed t-test. **e.** Tibia anterior muscles from WT and tumor bearing mice for 3 weeks were harvested and extracted total RNA to evaluate a hypoxic gene, *glut1* by RT-qPCR. Data are mean±SE for n=4-6 mice. The *p* values were evaluated by unpaired, two-tailed t-test. **f-g.** Mixed skeletal muscles (TA, GC, and quadriceps) from WT mice and melanoma bearing mice for 1-, 2-, and 3-weeks post tumor implantation (pti) were harvested and enzymatically digested. The inflammatory cells were stained with CD45 specific antibody by following FACS analysis. **f.** The inflammatory cells were represented by CD45+ cells of whole muscle cells. Gating strategy was presented at *Extended Data Fig.3b*. **g.** Quantification of CD45+ inflammatory cells in total muscle cells. The data was represented by percent of CD45+ cells of total muscle cells. Data are mean± SEM for n=4-5 mice. The statistical analysis was performed by one-way ANOVA, followed by post-hoc multiple comparison analysis using Tukey test, and the adjusted p-values were reported. **h.** Tibia anterior muscles from WT mice (5 month) and KPC mice (3, 4, and 5 month) were harvested and extracted total RNA to evaluate inflammatory genes, *ICAM1, IL-1b, IL-10,* and *IL-6* by RT-qPCR. Data are mean±SE for n=4-8 mice. The statistical analysis was performed by one-way ANOVA, followed by post-hoc multiple comparison analysis using Tukey test, and the adjusted p-values were reported.

Next, to determine whether the impairment of the endothelial barrier could also promote immune cell infiltration and inflammation in the muscles, autopsy-derived abdominal muscles from control and cancer cachexia patients were examined by H&E staining. As shown in **Extended Data Fig. 3a**, the muscles of cancer cachexia patients show high immune cell infiltration (dark violet color) near capillaries. To examine whether EC loss in cachexic muscle can trigger immune cell recruitment into muscles, the total number of immune cells (CD45+ cells) were determined in muscles of melanoma bearing mice in a time-dependent manner by flow cytometric analysis. The total number of CD45+ immune cells in muscle showed a major surge as early as week 1 and then gradually decreased over the course of tumor progression (**Figs. 3f-g** and **Extended Data Fig. 3b**). Furthermore, we found that the muscles of KPC-mice had increased expression of inflammatory mediators such as *ICAM1*, *IL-1β, IL-10*, and *IL-6* during cachexia progression (**Fig. 3h**).

### Unbiased analysis of dynamic differentially expressed genes in muscle vascular ECs during distal tumor growth

We next used unbiased transcriptomic analyses to elucidate the signaling pathways in muscle ECs that were affected by remote tumor growth. We first isolated skeletal muscle ECs from control and tumor bearing mice in a time dependent manner by flow cytometric sorting, and then performed bulk RNA-seq analysis. We focused on **D**ynamic **D**ifferentially **E**xpressed **G**enes (DDEG) in muscle vascular ECs during distal tumor growth using TrendCatcher^46^. We identified 2,180 DDEGs from 15,715 profiled genes, using an adjusted dynamic P value less than 0.05. We also identified top 10 up-regulated enriched biological pathways using DDEGs with positively accumulated log fold change over time, and together with top 10 down-regulated enriched biological pathways using DDEGs with negatively accumulated log fold change over time (**Extended Data Fig. 4a** and **Supplementary table 1**). To show the selected biological pathways GO enrichment change over time, we used the TimeHeatmap visualization function from TrendCatcher (**Fig. 4a**, *see details in the Method Section*). Pathway enrichment was highly dynamic the week following distal tumor implantation suggesting a strong early endothelial response to circulating cues released by the tumor. We found that 78 DDEGs which are associated with muscle cell differentiation were upregulated within one week with an averaged log2FC of 2.02. Of note, several of these genes such as *Hey1, Ski, Tbx1, CCn3, CCn4, Npnt, Myh11, Smad6, Edn1, Acta1, Actc1, Tnnt3, Myoz1,* and *Myoz2* are regulators of muscle cell differentiation and are typically upregulated in EndMT (**Fig. 4b**) which is consistent with our observation of increased α-SMA in muscle ECs (**Figs. 2g-h**). Moreover, we found that muscle vascular ECs dynamically increase expression of genes which are associated with energy metabolism (oxidative phosphorylation, ATP metabolic process) and reactive oxygen species (ROS) during distal tumor growth (**Fig. 4a**) but decrease expression of genes related to catabolic processes (**Extended Data Fig. 4b**). These data suggest that the remaining muscle ECs may compensate for the loss of EC function even though the vascular density is decreased during cachexia progression. We further analyzed individual genes which are associated with endothelial ROS metabolism. Interestingly, gene expression of anti-oxidant enzymes such as *Prdx2, Prdx4, Prdx5, Gpx1,* or *Sod3* was significantly induced in muscle ECs during distal tumor growth (**Extended Data Fig. 4c**) which may be related to hypoxia in cachexia muscle. The unbiased RNA-seq analysis using skeletal muscle ECs strongly supported the hypothesis that circulating factors from the tumor or tumor microenvironment remotely regulate expression of genes in skeletal muscle capillary ECs.

**Figure 4.**
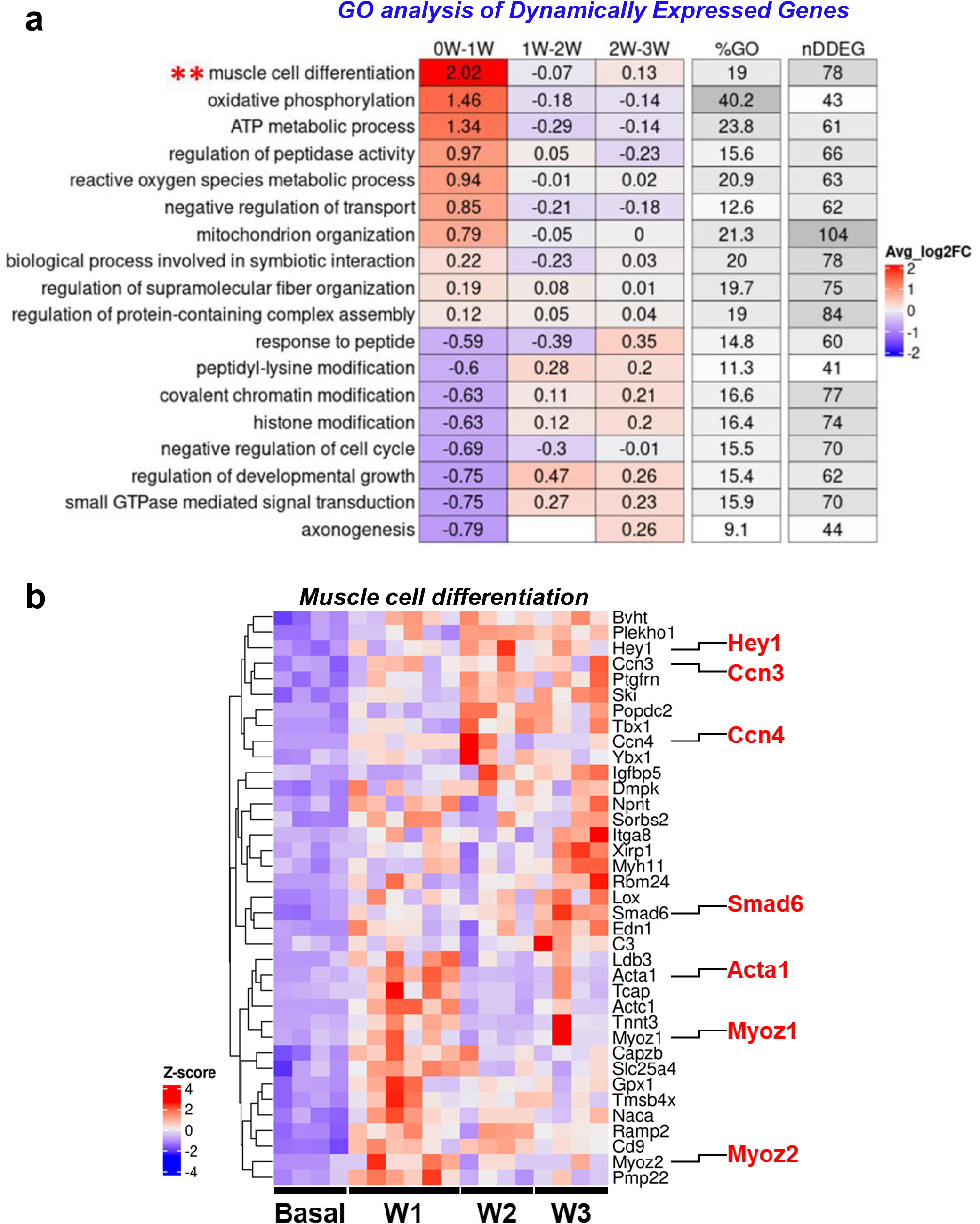
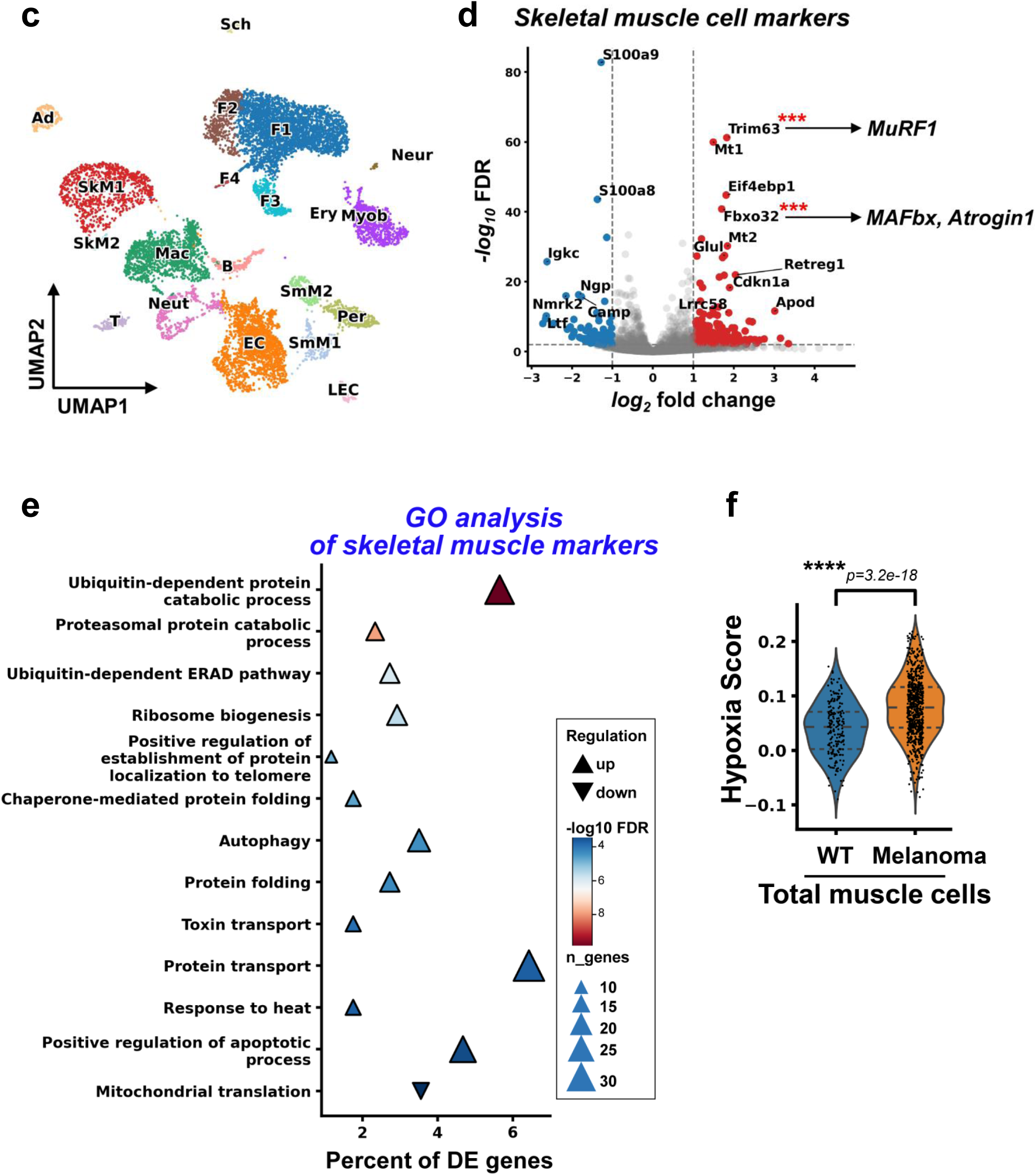
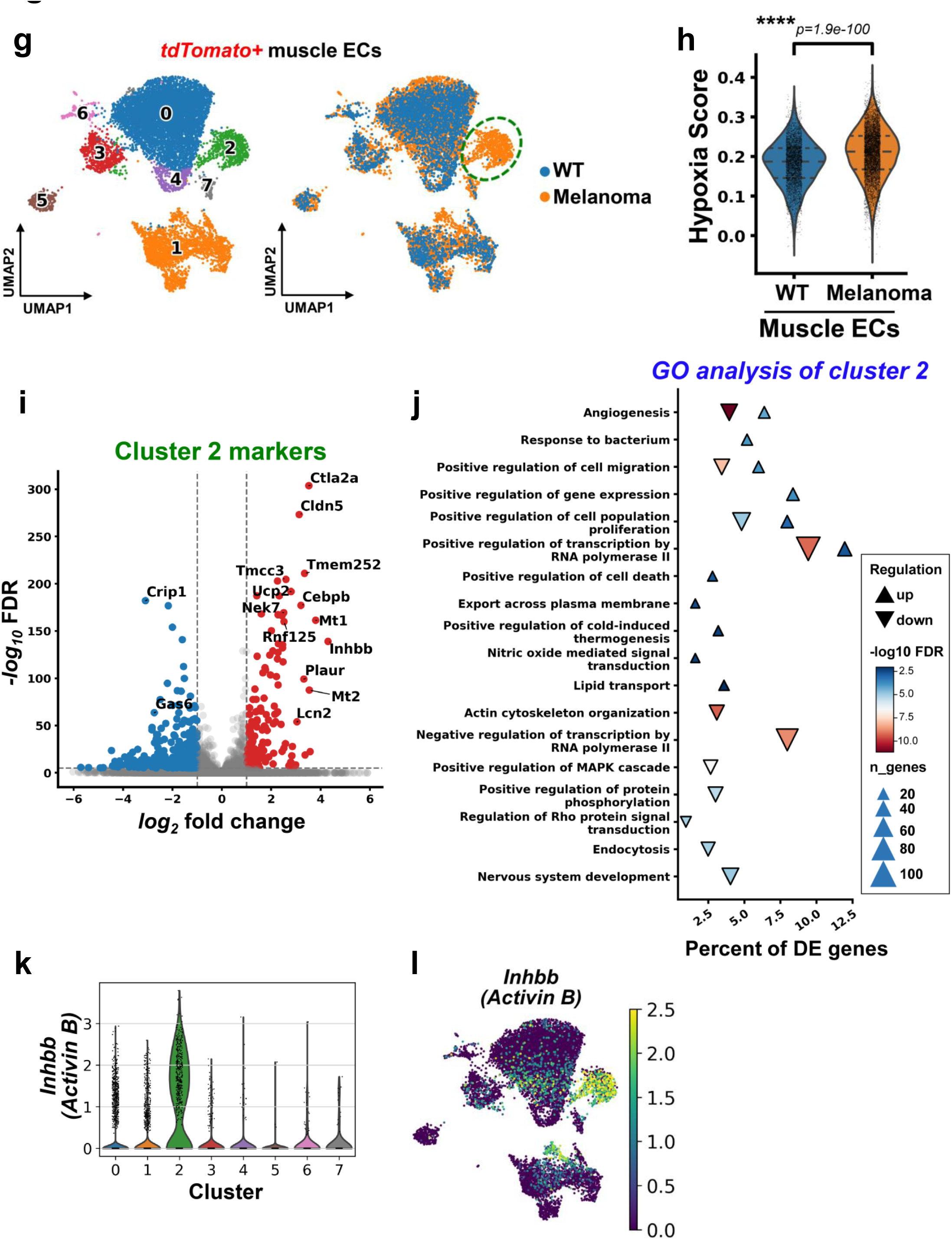
Dynamic differentially expressed genes and a specific EC subpopulation in endothelium of cancer cachexic muscles. **a-b.** Unbiased RNA-seq analysis of dynamic differentially expressed genes (DDEGs) in muscle vascular ECs during distal tumor growth. The mixed skeletal muscle ECs were isolated from TA, GC and quadriceps in WT and melanoma bearing mice in a time dependent manner (1-, 2-, or 3- weeks) and then subjected to bulk RNA-seq analysis. **a.** TimeHeatmap of the top dynamic pathways. Each column represents a time interval. “0W-1W” represents week 0 to week 1. The “%GO” column represents the percentage of DDEGs identified from the corresponding GO term. The “nDDEG” column represents the number of DDEGs from the corresponding pathway. The number within each cell represents the averaged log2 fold change (Avg_log2FC) of gene expressions compared to the previous time point. Color represents the magnitude of the change. For WT (n=4 mice), week 1 (n=6 mice), week 2 (n=4 mice), and week 3 (n=4 mice). **b.** DDEGs expression heatmap from the biological pathway of muscle cell differentiation **\*\*** in **a**. The genes (*Hey1, Ccn3, Ccn4, Smad6, Acta1, Myoz1*, and *Myoz2*) which are related with EndMT are highlighted. **c-i.** Single cell RNA-seq analysis of muscle cells and muscle ECs from WT or melanoma bearing mice. The skeletal muscle cells were isolated from TA, GC and quadriceps in tdTomato flf:Cdh5-CreERT2 (EC^tdTomato^) mice without (WT, n=3 mice) or with tumor (melanoma, n=4 mice) for 3 weeks. The muscle-ECs (tdTomato+DAPI-) and muscle-non-ECs (tdTomato-DAPI-) were collected by FACS sorting and subjected to scRNA-seq analysis. **c.** The integrated UMAP for tdTomato negative (-) mixed muscle cells from WT and melanoma bearing mice. The muscle cells show 20 total clusters and were assigned by cell specific markers. **d.** Volcano plot for differentially expressed skeletal muscle genes in melanoma bearing mice. **e.** GO analysis of differentially expressed skeletal muscle genes. **f.** Violin plot of hypoxia gene expression score in total muscle cells. **g.** The integrated UMAP of tdTomato positive (+) muscle ECs from WT and melanoma bearing mice. The muscle ECs show total 8 clusters (# 0-7, left panel) and cluster 2 is unique to melanoma bearing mice (dotted circle, right panel). **h.** Violin plot of hypoxia gene expression score in muscle ECs. **i.** Volcano plot for cluster 2 marker genes in melanoma bearing mice. **j.** GO analysis of cluster 2 marker genes. **k-l.** Inhbb (Activin B) expressing clusters of in violin plot (**k**) and integrated UMAP (**l**).

### Single cell transcriptomics identifies hypoxia, catabolism and inflammation signatures in endothelial subpopulations within the skeletal muscles of tumor bearing mice

Single-cell transcriptomics demonstrate that ECs have significant functional heterogeneity even within the same tissues^14, 47, 48^. Thus, we examined whether cachexic muscles have specific EC subpopulations that drive muscle loss using scRNA-seq analysis. We took advantage of inducible EC specific tdTomato lineage tracing mice (Cdh5-Cre^ERT2^-tdTomato, EC^tdTomato^)^49^ which would maintain their irreversible genetic endothelial lineage label with tdTomato. The skeletal muscles from the hindlimbs of tumor free EC^tdTomato^ mice (WT) and melanoma-cachexia EC^tdTomato^ mice were used to isolate the tdTomato positive (+) cells (ECs) and tdTomato negative (-) cells (non-ECs) by flow cytometric sorting and the isolated cells were subjected to scRNA-seq using the 10X Genomics 3’ platform.

First, we analyzed clusters of total muscle cells (tdTomato negative cells) from WT and melanoma-cachexia mice. The integrated mouse muscle cells clustered with 20 subpopulations and we assigned cell types by their expression of marker genes (**Fig. 4c** and **Supplementary table 2**). To evaluate genes differentially regulated in cachexic skeletal muscle, we compared differential expression between the skeletal muscle cells of melanoma bearing and healthy mice (**Fig. 4d**). The skeletal muscle cells from melanoma bearing mice had highly increased expression of cachexic marker genes such as *Trim63 (MuRF1)*, *Fbxo32 (MAFbx, Atrogin1)*, *Mt1/2*^50^ and *Retreg1* (autophagy regulator)^51^. GO enrichment analysis of the differentially expressed genes showed significant increases in catabolic processes through activating pathways of ubiquitination, proteasome, autophagy, protein transport, and apoptosis (**Fig. 4e**). Moreover, we determined whether the melanoma-induced cachexic muscles were subjected to hypoxia by scoring cells based on their expression of hypoxia response genes (GO:0071456). Skeletal muscle cells had a significant increase in hypoxia gene expression in melanoma-bearing mice compared to control mice (*p*=3.2×10^-18^) (**Fig. 4f**), as did most other cell types from the muscle tissue (**Extended Data Fig. 4d**). These results are consistent with our observations that KPC-cachexia mice have increased expression of cachexic marker genes (**Extended Data Fig. 1c**) and melanoma-cachexia mice show muscle vascular leakage and hypoxia (**Figs. 3a-e**).

Next, we analyzed whether cachexic muscles have specific EC subpopulations that may drive muscle loss. The integrated muscle ECs of WT and melanoma-cachexia mice clustered with seven subpopulations (the left UMAP plot of **Fig. 4g**). To determine whether muscle ECs also increase hypoxic response in cachexic muscles, we scored the ECs based on their expression of hypoxia response genes (GO:0071456). Muscle ECs (tdTomato positive cells) had a significant increase in hypoxia gene expression in melanoma-bearing mice compared to control mice (*p*=1.9×10^-100^) (**Fig. 4h**). Importantly, muscle ECs from melanoma-cachexia mice had a unique EC subpopulation, cluster 2, not present in control mice (**Fig. 4g**, the circled cluster in right panel). Therefore, we further analyzed differentially expressed genes in cluster 2 (**Fig. 4i** and **Supplementary Table 3**) and their functions with GO enrichment analysis (**Fig. 4j**). Angiogenesis was highly enriched in the genes downregulated in cluster 2 relative to other ECs, suggesting the unique population has reduced angiogenic potential and reduced EC functionality. The upregulated genes in cluster 2 were enriched for increased immune response signatures. We also found that expression of *Inhibb*, the gene which encodes for the β subunit of Activin and Inhibin, is the most differentially upregulated gene in cluster 2 muscle ECs (**Fig. 4k**). The gene encoding for the α subunit of Inhibin was not differentially regulated in cluster 2, suggesting the upregulation of Inhibb is specific to *Activin-B*. We further showed that *Inhibb* (*Activin-B*) is predominantly expressed in cluster 2 and only minimally expressed in other clusters (**Fig. 4l**). These scRNA-seq data indicate that a specific cluster 2 of skeletal muscle ECs shows inhibition of canonical endothelial function and increasing immunogenicity, thus possibly contributing to muscle atrophy.

### Endothelial PGC1α is critical for maintaining muscle homeostasis

We hypothesized that the observed changes in transcriptomic profiles were the result of tumor induced shifts in the activity of transcription factors. Therefore, we examined whether transcription factors were differentially active in the muscle ECs of melanoma-bearing mice by analyzing the promoter regions of differentially expressed genes for binding motif enrichment. We observed multiple enriched binding motifs (**Supplementary table 4**), for example our analysis found KLF15 an important transcription factor for endothelial homeostasis^52^ to be the most highly enriched binding motif in downregulated genes. Interestingly, the downregulated genes in cluster 2 were enriched (Fisher’s exact, BH corrected p < 0.05) for multiple cofactors of the transcriptional coactivator PGC1α such as Nrf1, PPARδ, Mef2, Hnf4, and Foxo1 (**Supplementary table 4**). Therefore, we interrogated a previously reported PGC1α ChIP-seq dataset^53^ and identified which genes were potentially regulated by PGC1α by their proximity to significant peaks. Downregulated cluster 2 genes were highly enriched for genes within 5000 bp of PGC1α peaks (Hypergeometric, *p*=2.4×10^-7^) and upregulated genes were mildly enriched (*p*=0.0002) (**Extended Data Fig. 4e**), likely representing the dual repressor and activating roles of PGC1α. Together, these data suggest that PGC1α is dysregulated in the muscle endothelial cells of tumor bearing mice.

PGC1α supports cell survival in many cell types by regulating energy and redox signaling^24^. Thus, we first examined whether expression of PGC1α was altered in skeletal muscle ECs from KPC-cachexia mice. Surprisingly, the muscle ECs of KPC-cachexia mice had significantly decreased protein levels of PGC1α compared to age-matched control mice (**Figs. 5a-b**). Moreover, the protein levels of PGC1α were consistently decreased in muscle ECs from melanoma bearing mice at week 1 and 2 after tumor implantation compared to control mice (**Extended Data Figs. 5a-b**). Next, we examined whether the increased circulating levels of Activin-A we had observed in cachexic mice could reduce PGC1α expression in muscle ECs. Interestingly, we observed Activin-A exposure decreased PGC1α protein levels (**Fig. 5c**) and PGC1α transcription (**Fig. 5d**) in ECs. Moreover, we determined whether Activin-A negatively regulates PGC1α promoter activity using PGC1α promoter luciferase plasmid. As shown in **Fig. 5e**, Activin-A treated ECs suppressed PGC1α promoter activity compared to TNFα treated positive control. Thus, we examined the effects of loss of EC-PGC1α in EC survival, EndMT, and vascular barrier function. PGC1α depleted ECs by lentivirus expressing PGC1α-specific shRNA (**Extended Data Figs. 5c-d**) showed significantly increased apoptosis relative to control cells (**Fig. 5f**). Conversely, the overexpression of EC-PGC1α in PGC1α depleted ECs restored the expression of anti-apoptotic gene, *Bcl2* (**Extended Data Fig. 5e**). PGC1α depleted ECs also significantly increased mRNA levels of the EndMT marker, *Vimentin* (**Fig. 5g**). The adherens junctional protein VE-cadherin is critical for the maintenance of the vascular barrier, and its breakdown leads to EC barrier permeability and increases pathology^54, 55^. Therefore, we next examined the role of PGC1α in regulating the adherens junctions of ECs by immunofluorescence (IF) staining for VE-Cadherin. EC-PGC1α depletion markedly disrupted endothelial barrier adherens junctions (**Fig. 5h**) and significantly increased mRNA levels of proinflammatory cytokine genes, *VCAM1, Caspase1*, and *IL-6* relative to control cells (**Fig. 5i**). Next, to determine whether PGC1α regulates VE-cadherin expression in ECs, we searched for putative binding motifs of PGC1α on the human VE-cadherin (*Cdh5*) promoter. However, the VE-cadherin promoter has two putative PPARγ binding motifs instead of PGC1α direct binding sites (**Extended Data Fig. 5f**). Thus, we designed specific primers to recognize two PPARγ binding motifs in a chromatin immunoprecipitation (ChIP) assay to determine whether PGC1α can indirectly bind PPARγ binding motifs on VE-cadherin promoter. The nuclear fraction was extracted from control and Activin-A treated ECs and the DNA was sheared by sonication and followed by immunoprecipitation with a PGC1α specific antibody. The pull-downed DNA was evaluated by qPCR with specific primers which recognized putative PPARγ binding sites on the VE-cadherin promoter. The anti-PGC1α antibody highly bound the VE-cadherin promoter in control ECs but is significantly inhibited by Activin-A treatment (**Fig. 5j**). Moreover, Activin-A treated ECs had significantly decreased protein levels of the adherens junction protein VE-cadherin (**Extended Data Fig. 5g**). Taken together, these results suggest that EC-PGC1α is required for EC survival and regulates expression of VE-cadherin by binding to the VE-cadherin promoter, and that PGC1α downregulation in the endothelium by Activin-A in cancer cachexia causes loss of muscle vascular barrier integrity.

**Figure 5.**
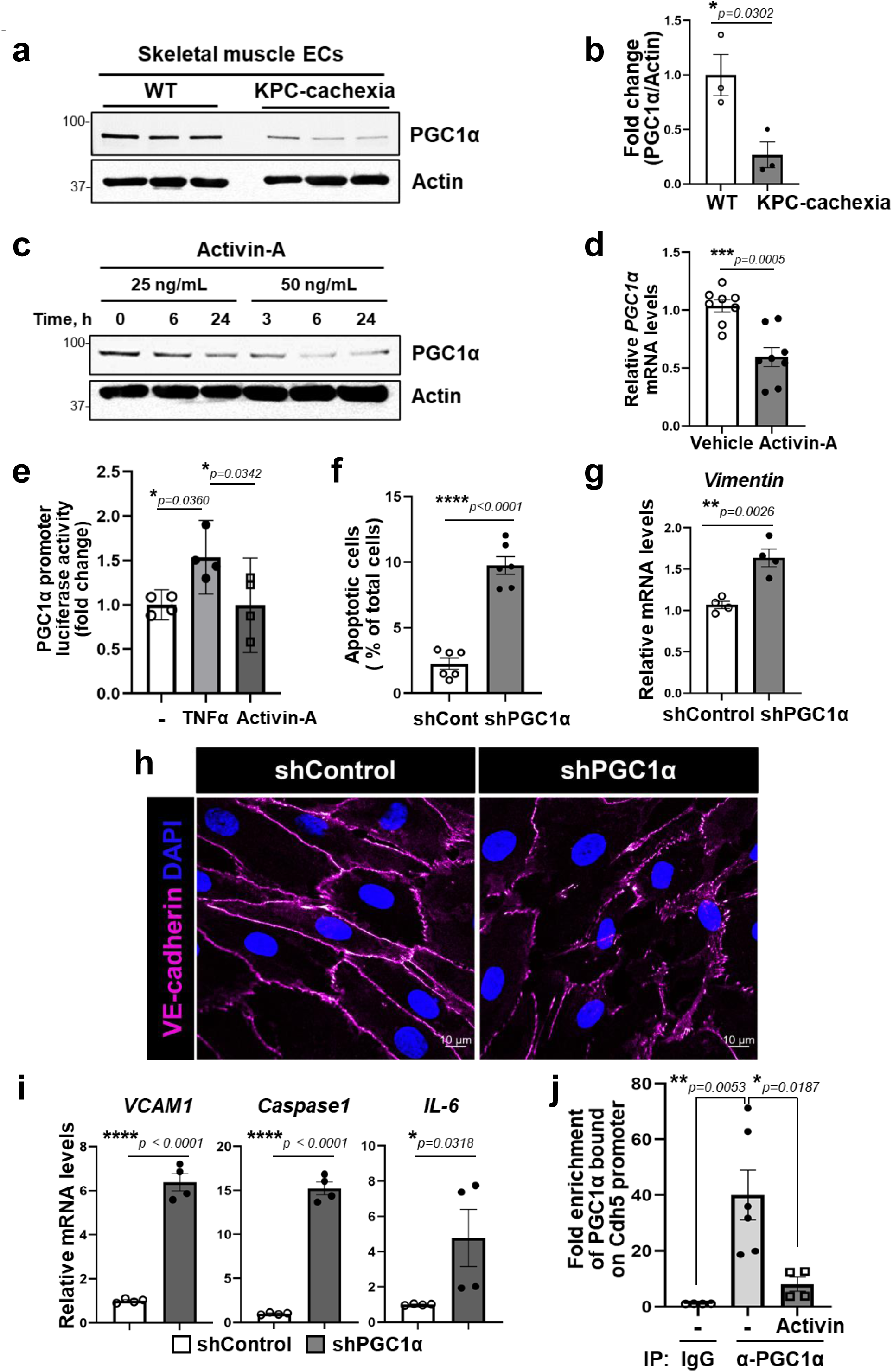
Circulating Activin-A induces dysfunctional vasculature by suppressing endothelial PGC1α in muscle. **a**. WT mice and KPC-cachexia (5 month) mice were harvested skeletal muscles (GC, TA, and quadriceps) from hind limbs. The mixed muscles were enzymatically digested and isolated muscle ECs using CD31-conjugated microbeads. PGC1α expression was determined by Western blotting with PGC1α specific antibody. Each lane represents one mouse. **b.** The levels of PGC1α in **a** were normalized by actin and presented fold change. Data are mean±SE for n=3 mice. The *p* values were evaluated by unpaired, two-tailed t-test. The original blots can be found in Source data. **c.** HMVECs were treated with Activin-A (25 ng/mL and 50 ng/mL) for indicated times (0, 3, 6, or 24 h). The cell lysate was determined for PGC1α expression by Western blotting with PGC1α specific antibody. The representative blots are from 3 independent experiments. The original blots can be found in Source data. **d**. HMVECs were treated with Activin-A (25 ng/mL) for 24 h. The *PGC1α* expression was determined by RT-qPCR with specific primers. The levels of *PGC1α* were normalized by *18s* levels and presented fold change. Data are mean±SE for n=8 biological replicates. **e.** HMVECs were transfected with PGC1α promoter luciferase plasmid and renilla for 48h and treated with Vehicle (0.1% BSA), TNFα (10 ng/mL), or Activin-A (25 ng/mL) for 16 h. The luciferase activity was normalized by renilla and presented fold change. Data are mean±SE for n=4 biologically independent samples. The statistical analysis was performed by one-way ANOVA, followed by post-hoc multiple comparison analysis using Tukey test, and the adjusted p-values were reported. **f-i**. HMVECs were treated with lenti-viral shRNA for Control and PGC1α for 72 h. **f**. The cell apoptosis was determined by staining with Annexin V-FITC and PI following FACS analysis. The apoptotic cells (Annexin V-FITC+PI+) were presented with percent (%) of total cells. Data are mean±SE for n=6 biological replicates. The *p* values were evaluated by unpaired, two-tailed t-test. **g.** The mRNA levels of *Vimentin,* a marker of EndMT were determined by RT-qPCR with their specific primers and normalized by *18s* levels and presented fold change. Data are mean±SE for n=4 biological replicates. **h.** EC barrier integrity was determined by IF staining with VE-cadherin antibody (magenta). Nucleus was stained with DAPI. The images were taken using confocal microscopy (LSM880, 60x objective) and represented from n=3 biological replicates. **i.** The mRNA levels of inflammatory markers, *VCAM1, Caspase1,* and *IL-6* were determined by RT-qPCR with their specific primers and normalized by *18s* levels and presented fold change. Data are mean±SE for n=4 biological replicates. **j.** The control and Activin-A treated ECs were crosslinked with 1% PFA for 10 min at RT and isolated nuclear fraction to shear DNA by following Chip assay with PGC1α specific antibody. Normal mouse IgG and anti-RNA polymerase antibodies were used for negative control and positive control, respectively. The pulldown DNA was amplified by qPCR with *Cdh5 (VE-cadherin)* promoter specific primers. The enrichment values were normalized with input values and presented fold changed. Data are mean±SE for n=4-6 biological replicates. The statistical analysis was performed by one-way ANOVA, followed by post-hoc multiple comparison analysis using Tukey test, and the adjusted p-values were reported.

### Overexpression of EC-PGC1α preserves muscle vascular density and prevents cancer cachexia

Next, we examined whether restoring EC function can prevent muscle loss in cancer cachexia. To evaluate EC specific effects of PGC1α in mouse muscle, we generated a lenti-viral construct (lenti- *<u>Cdh5</u>* -mPGC1α- *<u>cm v</u>* -GFP virus, hereafter referred to as lenti-EC-PGC1α-GFP virus) which co-encodes mouse PGC1α and GFP controlled by the EC specific promoter VE-cadherin (*Cdh5*) and by CMV promoter, respectively. After tumor implantation, the mice were intramuscularly injected with lenti-EC-PGC1α-GFP virus in five regions of the gastrocnemius and tibialis anterior muscles. The lenti-EC-PGC1α-GFP virus was specifically expressed in the muscle endothelium as confirmed by the colocalization of the EC specific marker CD31 and GFP (**Extended Data Fig. 6a**). EC-PGC1α overexpression preserved the muscle vascular density and CSA, which were decreased in tumor bearing mice (**Figs. 6a-c**). Moreover, EC-PGC1α overexpression showed a trend towards increased number of muscle fibers within the 1500-2500 µm^2^ range (**Extended Data Fig. 6b**). Mouse grip strength was also increased by intramuscular overexpression of EC-PGC1α in tumor bearing mice (**Fig. 6d**) and resulted in decreased mouse fatigue by recovering original force (**Fig. 6e**). Additionally, intramuscular overexpression of EC-PGC1α decreased expression of hypoxic marker gene, *Glut1* or cachexia marker genes, *MuRF1* and *Atrogin1* (**Fig. 6f**). Moreover, EC-PGC1α overexpression significantly inhibited expression of proinflammatory cytokine genes, *ICAM1*, *TNFα*, *IL-6*, *IL-10*, or *IL-1β* which were increased in tibialis anterior muscle of tumor bearing mice compared to control mice (**Fig. 6g**) even though mouse body weight and tumor volume were not significantly changed by EC-PGC1α overexpression (**Extended Data Figs. 6c-d**). Taken together, we propose that high levels of Activin in cancer remotely suppresses PGC1α expression in muscle vasculature which contributes to EC loss and vessel leakage, eventually causing muscle loss and cachexia. However, overexpression of EC-PGC1α protects ECs in muscle, normalizes EC function, and restores muscle mass, thereby preventing cachexia progression (**Fig. 6h**).

**Figure 6.**
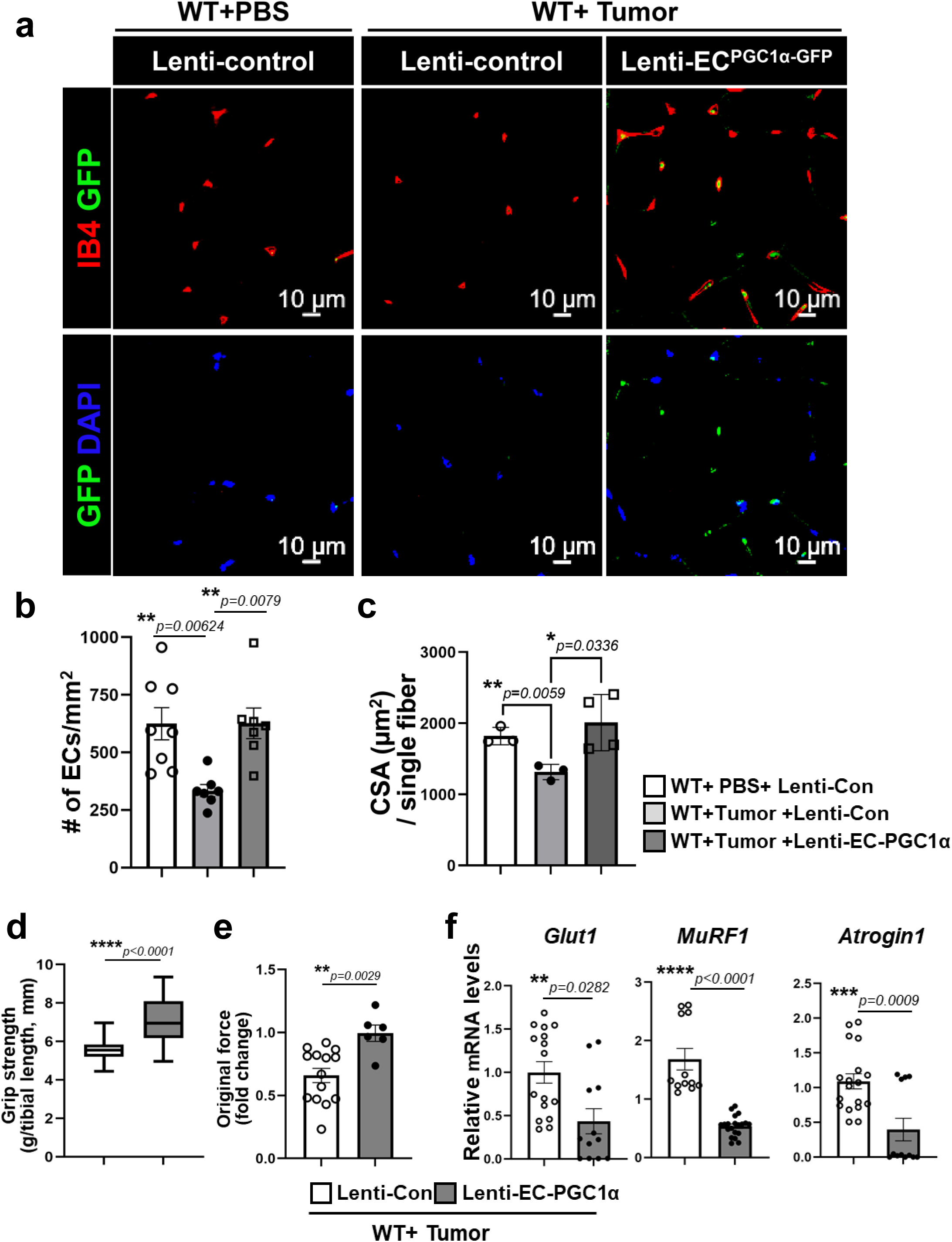
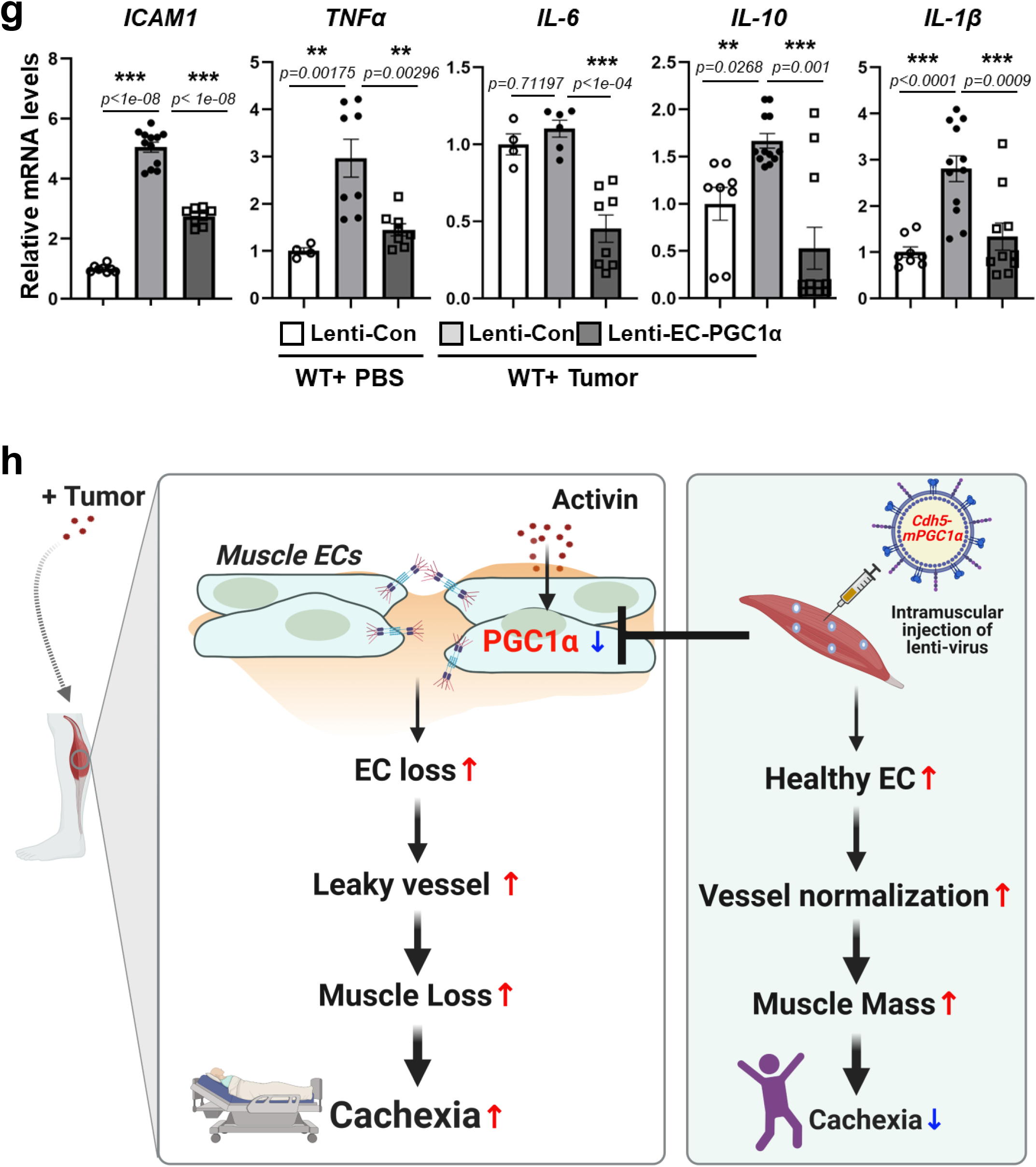
Overexpression of EC specific PGC1α in muscle of tumor bearing mice preserves vascular density and inhibits muscle atrophy. **a-c and g.** Melanomas were subcutaneously implanted in the dorsal flank of WT mice and then the mice were intramuscularly received with lenti-control and lenti-EC-PGC1α-GFP virus. Control mice were received with same volume of PBS instead of tumor and with lenti-control virus. All mice were examined in 3 weeks after tumor implantation. **a**. The cryosections of gastrocnemius were stained with EC-specific IB4 (red). The images were taken using confocal microscopy (LSM880, 20x objective) and representative images are from n=7-8 mice. Nuclei was stained with DAPI and exogenously overexpressed EC-PGC1α was evaluated by GFP fluorescence (green). **b.** Muscle vasculature density were determined by number of IB4+ ECs per field (mm^2^). Data are mean±SE for n=5-7 mice. The statistical analysis was performed by one-way ANOVA, followed by post-hoc multiple comparison analysis using Tukey test, and the adjusted p-values were reported. **c.** The cross-sectional area (CSA) of muscle was measured in single fiber using Image J. Data are mean±SE for n=3-4 mice. The statistical analysis was performed by one-way ANOVA, followed by post-hoc multiple comparison analysis using Tukey test, and the adjusted p-values were reported. **d-f.** All WT mice were subcutaneously implanted with melanomas in the dorsal flank and then the mice were intramuscularly received with lenti-control and lenti-EC-PGC1α-GFP virus. All mice were examined in 3 weeks after tumor implantation. **d.** The four limbs grip strength of mice was measured five times for each mouse using a grip strength test meter and were normalized by tibial length. Data are mean±SE for n=5-7 mice and 5 times measurements for each mouse. Data are presented with box and whiskers plot and whiskers are Min to Max. The *p* values were evaluated by unpaired, two-tailed t-test. **e.** The fatigue of mice was evaluated using the values of grip force in **d** and presented with fold change of decrease of original force. The *p* values were evaluated by unpaired, two-tailed t-test. **f.** Tibialis anterior muscles were analyzed expression of a hypoxia marker, *Glut1* or cachexia markers, *MuRF1* and *Atrogin1* by RT-qPCR, normalized by *PPIA* levels and presented with fold change. Data are mean±SE for n=12-18 mice. The *p* values were evaluated by unpaired, two-tailed t-test. **g.** The gastrocnemius from WT mice and tumor bearing mice with lenti-control or lenti-EC-PGC1α viruses were used to extract total RNA. The mRNA levels of proinflammatory genes (*VCAM1*, *TNFα, IL-6, IL-10,* or *IL-1β*) were determined by RT-qPCR, normalized by *PPIA* levels and presented with fold change. Data are mean±SE for n=4-12 mice. The statistical analysis was performed by one-way ANOVA, followed by post-hoc multiple comparison analysis using Tukey test, and the adjusted p-values were reported. **h.** *The proposed model;* health muscles (non-cachexia) have healthy capillary vessels and maintain muscle homeostasis, whereas high levels of Activin in cancer remotely suppress PGC1α expression in vasculature and the decrease of EC-PGC1α contributes EC dysfunction by increasing vessel leakage and causes muscle loss which is pivotal indicator for cachexia. However, overexpression of EC specific PGC1α in muscle can prevent EC loss, increase healthy ECs, contribute vessel normalization, and restore muscle mass and strength.

## DISCUSSION

Cachexia is not merely a side effect of cancer or cancer treatments. It needs to be treated with efficient clinical therapeutic approaches. Prior research has established the phenotypes of cachexia (muscle loss by increasing expression of E3 ubiquitin ligases, *MuRF1* or *Atrogin1*) and potential cachexia mediators including TNFα, IL-6, IL-1β, IFNγ, and TGFβ family members, which negatively impact muscle function^1,7, 56^. However, the role of vascular endothelial cells, which encounter circulating mediators before muscle cells do, has not yet been examined in cachexia. Therefore, we addressed fundamental questions such as: *Does the loss of muscle vascular density precede muscle loss in cancer cachexia? What are the circulating cues which induce muscle vascular dysfunction in cachexia? What are the roles of EC-PGC1α in muscle vasculature? Can targeted activation of pathways in the muscle endothelium prevent or reverse muscle loss in cachexia mouse models?*

In this study, we found that (1) cachexic muscles from tumor bearing mice or cancer patients have significantly decreased muscle vascular density which is associated with increase of vascular permeability and immune cell infiltration; (2) muscle ECs regulate muscle differentiation during cancer cachexia progression and a specific muscle EC subpopulation is associated with muscle atrophy; (3) in a cachexia model, circulating Activin-A suppresses expression of PGC1α in muscle vasculature, and induces vascular death and EndMT; (4) endothelial PGC1α regulates expression of the EC adherens junctional protein, VE-cadherin; and (5) increasing endothelial PGC1α expression prevents muscle loss in tumor bearing mice.

We first found that muscle endothelial dysfunction is a critical determinant of muscle loss in cachexia progression using two different mouse models: KPC mice, a genetic cancer cachexia model, and a melanoma allograft model. Traditional tissue histology is limited to small pieces of tissue and interpretation of thin section staining, thereby subject to multiple artifacts. However, we were able to perform comprehensive morphometric assessments of the muscle microvascular tree using a high-resolution 3D immunofluorescence microscopy, transparent tissue tomography (T3)^27, 28^. By quantitative analysis of 3D image data, we showed that cachexic muscles have impaired vasculature networks (**Fig. 1a**). Importantly, we observed that muscles of cancer cachexia patients also had significantly decreased muscle vascular density and muscle size (**Figs. 1h-k**). Similar findings were reported that the brains of SARS-CoV-2-infected patients have increased string vessels which represent lost capillaries^57^. Taken together, these suggest that intact and healthy capillaries play an important role to maintain cellular homeostasis in an organ specific manner. One of the underlying mechanisms of low muscle vascular density in cancer cachexia is increased cell death (apoptosis) as shown in **Figs. 2a-b**. These results suggest that increased apoptosis during cachexia progression results in vessel loss and decreased vascular network formation. Additional triggers for the loss of functional perfused vessels besides EC apoptosis, may be the loss of vasodilators or increased EndMT. TGFβ signaling is a known inducer of EndMT which can lead to a loss of endothelial function^58^. In recent years, the idea of the “*pre-metastatic niche*” has arisen which suggests that the tumor can affect vascular beds and tissues in several organs even before metastasis, for example, by increasing the permeability of the distant vasculature so that these tissues are “prepared” for the seeding of metastatic cells subsequently released by the tumor^59–62^. Moreover, cancer cachexia patients show paraneoplastic syndromes which accelerate the death of the host by manifesting at a distance from any tumor^63^. These strongly support possibility that the primary disease may remotely affect skeletal muscle vascular ECs, thus creating a ***“pre-cachectic niche”***. Importantly, we found that circulating Activin-A levels are significantly increased in the blood plasma of KPC-cachexia mice (**Fig. 2c**) and that Activin-A induced EndMT (**Figs. 2f-h**) may be associated with the vessel pruning in cachexia muscle. Mechanistically, the loss of vasculature in cancer cachexia mice causes muscle loss due to impaired perfusion, increased muscle capillary leakage, and muscle hypoxia which increases the expression of inflammatory cytokine genes and immune cell infiltration (**Fig. 3**). Our temporal analysis of immune cell infiltration into muscle during distal tumor growth shows high levels at early stages of tumor progression (1 week) which gradually subsided. Similar finding was recently reported that tumor remotely breaks down brain EC barrier and make a leaky brain^63, 64^. Taken together, our results suggest that the loss of functional muscle vasculature via apoptosis and EndMT as well as capillary leakage during the early phases of tumor progression create a local milieu of inflammation and microenvironmental hypoxia which in turn could be key triggers of muscle loss.

Unbiased transcriptomic analysis demonstrated an increased expression of muscle differentiation genes in skeletal muscle ECs of tumor bearing mice (**Figs. 4a-b**) despite of the fact that the tumors were not in proximity of the muscles. This suggests that muscle ECs remotely respond to circulating factors released by a distal tumor and undergo EndMT which both contributes to the decreased number of ECs as well as the impairment of key endothelial functions such as maintaining an intact endothelial barrier. Single cell transcriptomic analysis identified a unique muscle endothelial cell subpopulation (**Figs. 4g-l**) that was specifically enriched in tumor bearing mice and exhibited gene expression signatures of dysfunctional ECs such as decreased angiogenesis, proliferation, and migration along with increased expression of genes involved in cell death, ROS signaling, and enhanced immune responses. The most upregulated genes in this unique population were related to inflammation (*Ctla2a*, *Cd36*, *Cebpb*, *Rnf125*, *Ackr3*, *Ctla2b*), cell cycle (*Cdc42ep3*, *Nek7*, *Cdkn1a*), adipose tissue (*Ucp2*, *Lpl*, *Apold1*), and TGFβ signaling (*Inhbb*). Notably, *Inhbb* (*Activin-B*) was the most differentially upregulated gene. Our data suggests that circulating Activin-A from tumors or tumor microenvironments is one of the inducers of muscle EC dysfunction in cancer cachexia models (**Fig. 2c**). The significant expression of Activin-B in the melanoma-dependent muscle EC subpopulation (cluster 2) suggests that specific ECs are sensitive to circulating Activin-A and that this primed cluster 2 population further amplifies the negative autocrine effect of Activin-B on EC dysfunction. Activin is a 26 KDa dimer formed by two inhibin β subunits which have four variants βA, βB, βC, and βE. They bind to heteromeric tetramers consisting of two type 1 receptors (either ALK7/Acvr1c or ALK4/Acvr1b) and two type 2 receptors (Acvr2a or Acvr2b)^65^. Activin-A (βAβA) is a homodimer of inhibin βA and Activin-B (βAβB) is a heterodimer of inhibin βA and inhibin βB^66^. Taken together, we hypothesize that high levels of circulating Activin-A released by tumor cells can induce dysfunction of muscle ECs as manifested by impaired endothelial barrier integrity as well as to specifically increase an EC subpopulation expressing high levels of Activin-B which may exacerbate skeletal muscle EC dysfunction via autocrine or paracrine effects. The uniquely identified muscle EC subpopulation may represent a potential therapeutic target to prevent muscle atrophy in cancer cachexia.

We additionally found that Activin-A inhibits expression of EC-PGC1α at the transcriptional level in ECs, *in vitro* and *in vivo*. The depletion of EC-PGC1α increases cell apoptosis and increases EC barrier permeability by downregulating VE-cadherin, a key EC adherens junctional protein. ChIP assays showed that PGC1α induces expression of VE-cadherin by binding its promoter and this interaction is significantly inhibited by Activin-A (**Fig. 5j**). Our results suggest that EC-PGC1α is a critical regulator of EC barrier function and that loss of EC-PGC1α increases dysfunctional vessels in muscle and causes loss of muscle mass and function.

There also has been a shift away from traditional anti-angiogenic therapies towards tumor vessel normalization approaches^67^. Interestingly, vessel normalization improves vessel function^19, 68^, increases anti-cancer drug delivery^13^, promotes immunotherapy efficacy^69^ and complements traditional anti-angiogenic approaches. Importantly, we found that overexpression of EC-PGC1α specifically in the muscle endothelium was able to rescue grip strength and muscle mass in tumor bearing mice, thus providing a path for novel therapeutic interventions in which restoration of muscle endothelial health may be a prerequisite for restoring muscle health and function in cancer cachexia.

Recent reports suggest that the relative abundance of muscle fiber types is linked to some diseases such as type 2 diabetes^70^ and more muscle ECs lie next to oxidative fibers than glycolytic fibers^71^. We mainly evaluated tibialis anterior and gastrocnemius muscles which consist of mixed fibers (type I and II) in our cachexia mouse models. Thus, it is possible that the reduced muscle mass by loss of muscle ECs in cancer cachexia mice may be also associated with change of fiber type composition. It would be valuable to compare whether specific muscle types vary in their susceptibility to EC-PGC1α depletion. Moreover, it remains to understand whether the muscle ECs rescued by PGC1α overexpression can activate muscle regeneration or differentiation in cachexic conditions. It is also worth considering that intravenous infusion of lenti-EC-PGC1α virus may mimic an exercise effect in cachexic mice^72, 73^ or induce systemic vessel normalization which can prevent cachexia as well as tumor growth and metastasis^61^.

It remains to be seen whether the profound improvement in muscle mass and strength we observed by restoring muscle endothelial health via EC-PGC1α overexpression would translate to the human setting because clinical assessments of muscle strength encompass far more than grip strength or other measures typically assessed in animal models^74–76^. Interestingly, abdominal autopsied muscles from cancer cachexia patients also showed severe decrease in muscle vascular density compared to muscles from non-cachexia patients. These results highlight that the translational potential of our findings. We used female and male mice to generalize the roles of muscle vasculature in cancer cachexia but the sex differences in patients may be even more profound and require further evaluation of the therapeutic potential in sex-specific manner^44, 77^.

In summary, our findings establish the importance of skeletal muscle endothelial dysfunction as a key early pathogenic mediator in cancer cachexia and could serve as an important therapeutic target to prevent or reverse cachexia in cancer patients.

## Methods

### Mice

All animal studies were carried out following protocols approved by the Animal Care and Institutional Biosafety Committee of the University of Illinois at Chicago. Mice were maintained under standard conditions (standard diet and water) at 23 °C and ∼60% humidity with 12 h light and 12 h dark cycles. We purchased C57/BL6J mice from Jackson Laboratory (#000664) as control wild type (WT) mice. KPC (LSL-**K**ras^G12D/+^: LSL-Tr**p**53^R172H/R172H^: Pdx1-**C**re) mice were generated by crossing LSL-Kras^G12D/+^:LSL-Trp53^R172H/R172H^ mice with Pdx1-Cre homozygous mice^34^. We received KPC mice from Dr. Paul Grippo at University of Illinois at Chicago. Both sexes of mice were used for all experiments.

For tumor syngeneic allograft assay, mouse skin melanoma B16F10 cells (ATCC CRL-6475, 10^6^ cells/100 µL PBS) were implanted subcutaneously in the dorsal flank of WT mice (C57BL/6J). Control mice were injected with PBS. The mice were monitored every other day to measure tumor growth with electronic caliper and body weight for 3 weeks and tumor size was presented with tumor volume which was calculated with equation of width^2^ x height x 0.523 as previously described^78^. For lineage tracing of muscle endothelial cells during distal tumor growth, we used inducible endothelial specific tdTomato expressing mice (tdTomato flfl:Cdh5-CreERT2, EC^tdTomato^)^49^. To induce the Cre recombinase in tdTomato fl/fl: Cdh5-CreERT2 mice, 8 weeks EC^tdTomato^ mice were intraperitoneally administrated with tamoxifen (100 mg/kg, body weight, Sigma-Aldrich T5648) once a day for consecutive three days. One month later, the mice were used for tumor syngeneic allograft assay.

For rescue experiments of overexpression of EC specific PGC1α, we generated lentivirus expressing mouse PGC1α under endothelial specific Cdh5 (*CD144*) promoter. This lenti-Cdh5-mPGC1α-EGFP virus specifically express mPGC1α-EGFP in vascular ECs. Thus, the mice were injected intramuscularly with total 200 µL lentivirus (1×10^9^ pfu/µL, 40 µL per injection site) at 5 different areas of tibialis anterior and gastrocnemius right after tumor implantation. Lentivirus expressing empty vector were used as a control. All mice were examined in 3 weeks after tumor implantation.

### Mouse grip force and fatigue

The 4 limbs grip strength of mice was measured five times for each mouse using a Bio-GS3 Grip Strength Test Meter^5^ and the data are presented as averages after normalization against tibial length, which is not altered by muscle loss^79, 80^. The fatigue of mice was evaluated using the values of grip force. The degree of fatigue was estimated by comparing first two pulls with the last two pulls (formula (4+5)/(1+2) which would theoretically be 1 in mice that show no fatigue)^81^ and presented with fold change of decrease of original force.

### Mouse skeletal muscle collection

All mice were measured body weight and tibial length before dissecting tissues with digital caliper. Skeletal muscles (gastrocnemius, tibialis anterior, or quadriceps) from both hindlimbs were dissected and immediately weighed using a microelectronic weighing scale (Accuris instruments, W3100A-120) and then directly embedded in tissue plus O.C.T. compound (Fisher) on dry ice for cryosection. For paraffin blocking, the tissues were fixed with 10% neutral buffered formalin overnight and changed with 70% ethanol following paraffin embedding. The muscle mass (weight, g) was normalized with tibial length (mm). For Western blotting or RT-qPCR, the muscles were immediately frozen in liquid nitrogen and stored at −80 °C until use.

### Muscle clearing and 3D imaging

Gastrocnemius muscles were freshly harvested and fixed with 2% PFA for 10 min at RT. After washing in PBS, the muscles were longitudinally sectioned to 400 µm thickness using a vibratome (Leica, VT1200S) and stained with primary rat anti-mouse CD31 antibody (BioLegend, 102502, 1:1000 dilution) for 18 h at 4 °C. Then, the tissues were incubated with fluorescent secondary anti-rat IgG2 antibody at a 1:100 dilution for 18 h at 4 °C. The secondary antibody was prepared by conjugating anti-rat IgG2 antibody (BioLegend, 407502) with DyLight 633 (Thermofisher, 46414) at a 1:10 ratio for 18 h at 4 °C. For optical tissue cleaning, the muscle tissues were incubated in a gradient D-fructose solution (20% for 30 min, 80% for 30 min, then 100% for 1 h) at room temperature (RT). The 3D images were taken by using a confocal fluorescence microscope (Caliber ID, RS-G4) with a 40x oil objective (Olympus UPLXAPO 40XO, NA 1.4, 0.13 mm WD). The light source was 640 nm excitation (Toptica iChrome MLE-LFA 50 mW diode laser) with a 630/69 nm filter (Semrock) for DyLight 633. The images were reconstructed for visualization with Imaris Viewer 64-bit version 9.6.0. and the surface volume of vessels was analyzed with Imaris 64-bit version 7.2.2.

### Mouse skeletal muscle endothelial cell isolation

Skeletal muscles (gastrocnemius, tibialis anterior, or quadriceps) from hindlimbs were dissected, minced, and digested with digestion buffer (2 mg/mL collagenase A, 1 mg/mL dispase II, 0.1 mg/mL DNase I in 1x PBS without Ca^2+^/Mg^2+^) at 37 °C for 30 min with gentle shaking. At the end of the digestion process, the tissue was titrated using 18 G needles in syringes up and down 5 times and the cell suspension was filtered through 100 µm disposable cell strainer into a fresh 50 mL tube and centrifuged at 300x g for 5 min at 4 °C. The cell pellet was resuspended with wash buffer (20% FBS, 2 mM EDTA in 1x PBS without Ca^2+^/Mg^2+^) and filtered through 70 µm disposable cell strainer into a fresh 50 mL tube and centrifuged at 300 xg for 5 min 4 °C. This step was repeated with 40 µm disposable cell strainer into a fresh 50 mL tube. The cells were then resuspended with antibody binding buffer (0.5% BSA, 2 mM EDTA, 1% FBS in 1x PBS without Ca^2+^/Mg^2+^). To evaluate relative number of muscle EC in total muscle cells, the same number of cells (10^6^ cells/100 µL) were stained with specific antibodies of CD31 (EC marker, eBiosciences 14-0311082) and CD45 (immune cell marker, BioLegend 103108) for 30 min on ice. Samples were run through a Gallios flow cytometer (Beckman Coulter, Pasadena, CA) and analyzed by Kaluza software (Beckman Coulter). Because some immune cells also express CD31, leukocytes and dead cells were excluded by CD45+ and DAPI+ gating and the CD31+CD45-DAPI-cells were considered as pure ECs and represented by the percent (%) of total muscle cells. The muscle ECs (CD31+CD45-DAPI-) from single cell suspension of total muscle cells were obtained by FACS sorting using MoFlo Astrios (Beckman Coulter) or also isolated using mouse CD31 microbeads (130-097-418, Miltenyi Biotec) and MS column (130-042-201, Miltenyi Biotec) with magnetic separator for some experiments.

### Immunofluorescence imaging using confocal microscopy

*<u>1. Muscle tissue imaging for</u> <u>cryosection</u>-* The 7 µm cross-cryosections of muscle were dried for 5 min at RT and washed with 1x PBS for 5 min. The sections were blocked with blocking buffer (1x PBS, 2% BSA, 0.05% Tween 20, and 5% goat serum) for 1h at RT and then stained with EC specific CD31 antibody (BD biosciences, 550274, 1:25 dilution) or muscle base membrane laminin antibody (Sigma, L9393, 1:100 dilution) overnight at 4 °C in a humidified chamber. The secondary antibodies were used with goat anti-Rat IgM cross-adsorbed secondary antibody DyLight 488 (Thermo fisher, SA5-10010, 1:400 dilution) or goat anti-Rabbit IgG (H+L) cross-adsorbed secondary antibody Alexa fluor 633 (Thermo fisher, A21070, 1:500 dilution) for 1h at RT, respectively. The nuclei were stained with DAPI. *<u>2. Muscle tissue imaging for paraffin section</u>-* The 4 µm cross-paraffin sections of muscle were deparaffinized and hydrated at RT as following; 100% Xylene1, 5 min →100% Xylene2, 5 min →100% EtOH, 3 min →95% EtOH, 3 min →70% EtOH, 3 min →50% EtOH, 3 min → distilled water, 5 min → TBST (0.1% Tween 20), 5 min. For antigen retrieval, the slides were boiled in Sodium Citrate retrieval buffer (pH 6.0, 0.05% Tween 20) for 30 min and cooled for 20 min at RT. After washing with 1x PBS, the slides were permeabilized with 0.2 % Triton X-100 for 10 min at RT and followed by blocking with blocking buffer (1X PBS, 2% BSA, 0.05% Tween 20, and 5% goat serum) for 1 h at RT and then stained with EC specific biotinylated isolectin B4 (Vector lab, B-1205-.5, 1:100 dilution) overnight at 4 °C and followed by staining with FITC- streptavidin (Invitrogen, 11-4317-87, 1:400 dilution) for 1 h at RT. The images were taken by fluorescence microscopy or confocal microscopy (LSM880, x20 objective). *<u>3. In vitro cell imaging</u>-*The primary endothelial cells were cultured on coverslips in 6 well plates. After treatment with lenti-shRNA virus for indicated times, the cells were fixed with 4% PFA for 10 min at RT and then blocked with blocking buffer (1X PBS, 2% BSA, and 0.05% Tween 20) without permeabilization for 1 h. EC barrier integrity was visualized by immunostaining with 1:250 dilution of Alexa Fluor647 mouse anti-human CD144 (BD561567) for 1 h at RT. After mounting with antifade reagent with DAPI (Vector lab, H-1800-10), the images were taken using confocal microscopy (Zeiss LSM880, Plan Apo 1.46NA, 63x objective).

### Human abdominal muscle sections

We obtained human muscle sections from de-identified FFPE blocks from the Pathology archives through the UIC IRB-approved Pathology Biorepository. Data was not collected via interactions/interventions with individuals for research purposes, and there was no use of private, identifiable information about subjects. The paraffin sections (4 µm) were stained with H&E by UIC histology core laboratory. The images were taken using Olympus BX51/IX70 microscopy (x20 and x40 objectives). Human endothelial cells in muscles were stained with biotinylated Ulex Europaeus Agglutinin I Lectin (UEA I, B-1065-2, Vectorlab, 1:100 dilution) and FITC-streptavidin (11-4317-87, Invitrogen, 1:250 dilution). Nuclei was stained with DAPI. The images were taken using confocal microscopy (LSM880, x20 objective).

### Muscle capillary permeability assay

WT and tumor bearing mice for 3 weeks were anesthetized with Ketamine/xylazine and retro-orbitally administrated with FITC-albumin (250 mg/kg in PBS)^82^. After 10 min, circulating FITC-albumin was washed away by PBS perfusion and skeletal muscles (gastrocnemius and Tibial anterior) were immediately dissected and directly embedded in tissue plus O.C.T. compound on dry ice for cryosection. The frozen muscles were sectioned with 50 µm thickness using cryostat and dried for 5 min at RT, followed by PBS wash and directly mounted with antifade reagent with DAPI (Vector lab, H-1800-10). The FITC-albumin fluorescence images were taken by Z-sectioning using confocal microscopy (LSM710, X20 objective) and reconstructed for visualization using Imaris 64 bit 7.2.2. The intensity of FITC-albumin fluorescence in muscle was measured using Image J (1.52d, Java 1.8.0_172[64 bit]).

### Muscle hypoxia assay

WT and tumor bearing mice for 3 weeks were intraperitoneally injected with hypoxic probe pimonidazole (100 mg/kg, Hypoxyprobe, Hypoxyprobe plus kit) 1 h before harvesting muscle^83^. The paraffin muscle sections were used to measure the extent of hypoxia in gastrocnemius muscle by following the manufacturers’ guide. Nuclei were stained with DAPI and the images were taken using confocal microscopy (LSM880, X20 objective).

### *In situ* <u>T</u>erminal deoxynucleotidyl transferase d <u>U</u> TP <u>n</u>ick <u>e</u> nd labeling (TUNEL) assay

To determine apoptotic endothelial cells in muscles from KPC mice, the paraffin sections were deparaffinized, hydrated, and antigen retrieval was performed as described above. The apoptotic cells in sections were determined using TUNEL assay kit for *in situ* apoptosis detection (Invitrogen, C10618). After finishing TUNEL staining, the sections were stained with EC marker isolectin B4 (IB4) for overnight and followed FITC-streptavidin staining for 1h at RT. Nuclei were stained with DAPI and the images were taken by confocal microscopy (LSM880, x20 objective).

### FACS analysis for infiltrated immune cells in muscle

Briefly, mixed skeletal muscles (gastrocnemius, tibialis anterior, or quadriceps) from hindlimbs of control and tumor bearing mice were dissected, minced, and digested with digestion buffer (2 mg/mL collagenase A, 1 mg/mL dispase II, 0.1 mg/mL DNase I in 1x PBS) at 37 °C for 30 min with gentle shaking. Single cell suspensions were counted and the same number of cells (10^6^ cells/100 µL) were stained with 1:100 dilution of anti-CD45 (157610, BioLegend) antibody and DAPI. Samples were run through a Gallios flow cytometer (Beckman Coulter, Pasadena, CA) and analyzed by Kaluza software (Beckman Coulter). The inflammatory response (CD45+DAPI-) cells were represented by the percent (%) of total muscle cells.

### Blood plasma Activin level by ELISA

Mouse whole blood was slowly withdrawn by cardiac puncture and collected in anti-coagulant (0.5 M EDTA, 5 µL/100 µL blood) containing tubes. The blood was centrifuged in 2000x g at 4 °C for 15 min and plasma was collected in new tubes and stored at −80 °C until use. Activin-A levels in plasma were measured with 100 µL plasma for each sample using ELISA Kit (DAC00B).

### shRNA, plasmids, or lentivirus production

Five lenti-shPGC1α RNA constructs were purchased from Sigma (TRCN0000364084, TRCN0000364085, TRCN0000364086, TRCN0000001166, TRCN0000001167) and knockdown efficiency was examined in ECs. The TRCN0000001166 clone showed the best effect to deplete PGC1α in ECs and we used this clone for experiments. pcDNA4-myc-PGC-1α (Plasmid #10974) and PGC-1α promoter luciferase delta CRE (Plasmid #8888) were purchased from Addgene. pRL/TK (renilla-luciferase) was provided by Dr. Chinnaswamy Tiruppathi at University of Illinois at Chicago. Mouse endothelial specific lenti-viral vector for PGC1α (pLV-EGFP-Cd144_mPpargc1α [NM_008904.2]) were designed and produced by Vector builder. Lenti-viruses were produced in HEK293T cells (CRL-11268, ATCC) by transfection with DNAs (2.5 µg pMD2.G, 5 µg of psPAX2, and 7.5 µg of DNA expression vector) with 30 µg polyethylenimine (PEI, Polysciences, 23966, USA), and concentrated with Lenti-X concentrator (CloneTech, 631232) as described previously^84^.

### Cell culture

Human lung microvascular endothelial cells (HLMVECs, CC-2527, Lonza) were obtained from Lonza and cultured with EGM2 (Lonza) including all supplements and 10% FBS (Hyclone) until passage 8. The cells were transduced with lenti-shPGC1α RNA virus with 1:2000 dilution polybrene (Millipore, TR-1003-G) for 24 h and then media was changed. After 72 h from virus infection, the cells were used for experiments. For Activin treatment, the cells were starved with 2% FBS media overnight and treated with Activin-A (R&D system, 338-AC-010, 25 ng/mL or 50 ng/mL) for indicated times.

### Cell apoptosis

To evaluate cell apoptosis, the cells were stained with Annexin-V-FITC and PI (Bio-Rad, ANNEX20F) following FACS analysis using Gallios flow cytometer (Beckman Coulter, Pasadena, CA). The data was analyzed by Kaluza software (Beckman Coulter). The apoptotic cells (Annexin V-FITC+PI+) were presented with percent (%) of total cells.

### Western Blotting

The cells were lysed with lysis buffer (50 mM HEPES pH7.5, 120 mM NaCl, 5 mM EDTA, 10 mM Na pyrophosphate, 50 mM NaF, 1mM Na_3_VO_4,_ 1% Triton X-100). The concentration of protein was measured with Bradford protein assay solution (Bio-Red, 5000006) and the same amount of total protein was loaded in SDS-PAGE for Western blotting and probed with specific antibodies; PGC1α (NOVUS, NBP1-04676, 1:1000 dilution), Actin (Santa Cruz, sc-517582 HRP, 1:1000 dilution), or VE-cadherin (Cayman #160840, 1:1000 dilution). Quantitative analysis of Western blotting was performed using Image J (1.52d, Java 1.8.0_172[64 bit]). All raw gel blots are available in the Source data file.

### Quantitative real-time PCR

Total RNA was isolated by using TriZol Reagent (Invitrogen, 15596026). The muscle tissues were lysed with TriZol and homogenized in safe-locked 1.5 mL tubes with metal beads (Next advance, SSB14B) using tissue homogenizer (Next advance, bullet Blender, BBX24B-CE). Reverse transcription was carried out using high-capacity cDNA reverse transcription kit (Applied Biosystems, 4368814) using 1-2 µg of total RNA. Quantitative PCR for human genes (*PGC1α, Mcl1, Bcl2, Snail, Vimentin, VCAM1, Caspase1, IL6,* or *18S)* and mouse genes (*PGC1α, MuRF1, Atrogin1, Twist, Glut1, ICAM1, TNFα, IRF7, IL-10, IL-1*β*, IL-6,* or *PPIA)* was performed in duplicates with fast start universal SYBR Green master (ROX) PCR kit (Roche, 04913914001) using QuantStudio7 (Thermofisher). Expression of genes was normalized and expressed as fold-changes relative to *18S* or to *PPIA*. A complete list of all primers is available in the Source Data files.

### PGC1α promoter luciferase assay

ECs were transfected with 1 µg of a PGC1α promoter luciferase (Addgene # Plasmid #8888) and 35 ng of pRL/TK using PEI transfection reagent (Polyethylenimine). 48 h after transfection, the cells were stimulated with vehicle (0.1% BSA), TNFα (10 ng/mL), or Activin-A (25 ng/mL) for 16 h and then 100 µL of cell lysate from each sample was used to measure reporter gene expression. Firefly and Renilla luciferase activity were determined by the dual luciferase reagent assay system (Promega). The relative luciferase activity represents the mean value of the firefly/Renilla luciferase.

### Chromatin immunoprecipitation assay

Confluent ECs in 100 mm dishes were fixed with 1% PFA for 10 min at RT and washed with 1x cold PBS. To quench unreacted PFA, the cells were incubated with 1x Glycine for 5 min at RT and washed with 1x cold PBS. The nuclear fractions were obtained by using EZ-Magna Chip A/G kit (Millipore, #17-10086) and then followed by DNA shearing using Covaris (bath temperature 7.5 °C, acoustic power 3 W, duty cycle 2%, time 60 sec, cycle number 200). The sonicated nuclear fractions were centrifuged at 10k xg at 4 ⁰C for 10 min and then chromatin immunoprecipitation (ChIP) assay was followed with manufacturers’ guidance for EZ-Magna Chip A/G kit (Millipore, #17-10086) with minor modification. Each 50 µL aliquot was used for immunoprecipitation with 1 µg antibodies for normal IgG (negative control), RNA polymerase (positive control) or PGC1α antibody (NOVUS, NBP1-04676) for 3 h. For input control, 5 µL supernatant was aliquot and stored at 4 ⁰C until protein/DNA elution. The complex of protein and DNA (both input and immunoprecipitated samples) was incubated in elution buffer with proteinase K and RNase A at 37 ⁰C for 30 min and further incubated at 62 ⁰C overnight and then denatured at 95 ⁰C for 10 min. The purified DNA (2 µL) was enriched by qPCR (initial denaturation 94 ⁰C 10 min, 1 cycle; denature 94 ⁰C 20 sec, anneal and extension 60 ⁰C 1 min, 50 cycles) with primers for putative binding motifs of PPARγ on *VE-cadherin (Cdh5)* promoter. The gene amplified values were normalized with input values and presented with fold enrichment of normal IgG (negative control).

### Bulk RNA-seq and computational data analysis

ECs from skeletal muscles (gastrocnemius, tibialis anterior, or quadriceps) from hindlimbs were isolated and stained with specific antibodies of CD31 and CD45, and pure muscle ECs (CD31+CD45-DAPI-) were sorted using MoFlo Astrios (Beckman Coulter) as mentioned above. The sorted ECs were used for total RNA extraction by TRIzol Reagent (Invitrogen, 15596026) and the RNA was treated with DNase and its quality was evaluated with gel QC. Bulk RNA sequencing for 4-6 samples (each sample indicates a mouse) with 4 time points was performed with oligo-dT mRNA directional using Illumina Novaseq (HiSeq SR50) for coding genes at Genomic Facility in University of Chicago. Sequenced reads were aligned to the Mus musculus reference genome GRCm39 (mm39) with STAR v.2.7.6a^85^. Then mRNA expression counts were quantified from the aligned reads by using STAR *–quantMode* option. Genes were annotated using biomaRt R package^86^. We applied *calcNormFactors* function from edgeR R package^87^ to normalize the counts. Principal component analysis (PCA) was performed after normalization. Dynamic differentially expressed genes (DDEGs) were identified using TrendCatcher^46^ with adjusted dynamic p-value <0.05 threshold. For each DDEG, we calculated its accumulated log2 fold change (log2FC) over the time compared to its baseline expression. Then, we performed Gene Ontology (GO) enrichment analysis on both positively and negatively accumulated log2FC DDEGs respectively using clusterProfiler R package^88^, and picked the top 10% most enriched dynamic biological pathways. After removing redundant GO terms, for each biological pathway we calculated the averaged accumulated log2FC (GO_mean_logFC) from its corresponding DDEGs to infer the biological process accumulative change. Based on the pathway accumulative change ranking, we selected top 10 positively and negatively changed biological pathways respectively. To show how these GO enrichment change over time, we applied draw_TimeHeatmap_GO() function from TrendCatcher. To build the TimeHeatmap, TrendCatcher first check all the DDEG’s changing direction from its inferred trajectory within each time interval (either up or down). Then it calculated GO enrichment analysis for both up and down directed genes within that time interval. For each GO term within a corresponding time window, it also calculated a series of log2FC (Avg_log2FC) gene expression values over time, to quantify the trajectory dynamics of each biological pathway. Then we subset the TimeHeatmap object using draw_TimeHeatmap_selGO() function to show only the top 20 GO terms with the highest accumulative change.

### Single cell cDNA library preparation

EC^tdTomato^ mice with or without tumor for 3 weeks were used to dissect skeletal muscles (gastrocnemius, tibialis anterior, or quadriceps) from hindlimbs. The muscle tissues were minced and enzymatically digested with digestion buffer (2 mg/mL collagenase A, 1 mg/mL dispase II, 0.1 mg/mL DNase I in 1x PBS) at 37 °C for 30 min with gentle shaking as mentioned above. Control samples were pooled from 3 mice and melanoma samples were pooled from 4 mice. The single muscle cell suspension was sorted by tdTomato+DAPI-(ECs) and tdTomato-DAPI- (non-ECs) using MoFlo Astrios (Beckman Coulter). The sorted cells were loaded into a 10x Genomics microfluidics chip and encapsulated with barcoded oligo-dT– containing gel beads using the 10X Genomics Chromium controller according to the manufacturer’s instructions. Cell viability was more than 90% with a target of sequencing 6000 cells. Single-cell libraries were constructed using a Chromium Single Cell 3’ reagent kit v3.1 according to the manufacturer’s instructions. The quality of cDNA libraries was checked before sequencing and the cDNA libraries were multiplexed into one lane for sequencing on NovaSeq S1 (28 × 91bp paired-read).

### scRNA-seq data processing and analysis

Raw counts tables were generated for each sample with Cell Ranger (v6.0.2) using default parameters and the 10x mm10-2020-A reference. A total of 24107 cells passed quality control (**muscle ECs:** control tdTomato positive (+) 4046 cells, melanoma tdTomato positive (+) 7123 cells, **muscle non-ECs:** control tdTomato negative (-)5280 cells, melanoma tdTomato negative (-) 7658 cells). Data analysis was performed using the Scanpy (v1.9) python package^89^. Cells were filtered if they had mitochondrial read counts above 20% or if they expressed a genes-by-count ratio within the top 2% or bottom 2% of all cells. Data were normalized to 10,000 reads per cell using normalize_total() and the highly variable genes were found with highly_variable_genes(). The effects of total counts per cell and mitochondrial counts were regressed out with regress_out(). Each gene was scaled to unit variance and clipped at a value of 10 with the scale() function. Principle component analysis was performed using the pca() function on the variable genes. The neighborhood graph was computed using neighbors() with the top 20 principal components. Clustering was done with the Leiden algorithm then projected to two dimensions using umap(). Corresponding samples were integrated with the ingest() function. For TD tomato negative samples, cluster marker genes were found with rank_gene_groups() and cell types were annotated using well established markers in combination with markers from PangloaDB^90^. Differential expression testing between melanoma and control cells were performed with the DiffEXpy python package^91^ using the test.wald() function on the raw normalized counts. Multiple testing correction was performed using the Benjamini-Hochberg method. Differentially expressed genes and marker genes were tested for enrichment using the goatools^92^ python package and all musculus coding genes as the background. Statistically significant enrichments were defined by a corrected P value (Benjamini-Hochberg method) less than or equal to 0.05. The MEME Suite Simple Enrichment tool (https://www.biorxiv.org/content/10.1101/2021.08.23.457422v1) was used to search for JASPAR (https://doi.org/10.1093/nar/gkab1113) vertebrate transcription factor binding motif enrichment in the promoter regions 1000 bp upstream and 500 bp downstream of the transcription starting sites.

### Analysis of PGC1α binding enrichment

Previously reported ChIP-seq data from mouse adipose cells^53^ were used to annotate genes within 5000 bp of significant MACS peaks which intersected with study genes (n = 1112). Using a background of 15960 genes present in the study cells, the intersection of differentially downregulated and upregulated genes in EC cluster 2 were tested for overrepresentation using a hypergeometric test and visualized using GSEApy.

### Statistics and Reproducibility

Quantitative analysis of images and Western Blotting was performed using ImageJ (NIH, 1.52d, Java 1.8.0_172[64 bit]) software. Quantification is presented as the mean ± SEM from at least 3 independent biological replicate experiments or mice. The Student t-test with unpaired or paired tests were used for two group comparisons to determine statistical significance, with a p value threshold of less than 0.05. Significance levels are indicated in the figures as *p < 0.05, **p < 0.01, ***p < 0.001, and ****p < 0.0001. The t-test analyses were conducted using Prism 9.3.0, GraphPad Software (La Jolla, CA). We also used one-way ANOVA for statistical analysis using aov() function in R, followed by post-hoc multiple comparison analysis using Tukey test. Adjusted p-values were reported and less than 0.05 is considered as statistically significant. The significance levels of adjusted p values are indicated in the figures as *p < 0.05, **p < 0.01, and ***p < 0.001.

## DATA AVAILIBILITY

The authors declare that all data supporting the findings of this study are available within the paper and its supplementary information files. Source data files provided with this paper.

Bulk RNA-seq data is available at following NCBI GEO link: https://www.ncbi.nlm.nih.gov/geo/query/acc.cgi?acc=GSE211266

Single cell RNA-seq data is available at following NCBI GEO link: https://www.ncbi.nlm.nih.gov/geo/query/acc.cgi?acc=GSE211300

## CODE AVAILIBILITY

The TrendCatcher software platform used for longitudinal analysis of transcriptomic data is available at: https://github.com/jaleesr/TrendCatcher

## ACKNOWLEDGMENTS

The studies were supported by NIH grants P01-HL060678 (to JR), NIH grants P01-HL160469 (to JR), R33-CA258012 (to JR), T32-HL139439 (to MAS), R35-GM142743 (to SSYL), and AHA CDA grant 19CDA34680000 (to YMK). We were assisted by the Fluorescence Imaging Core at the Research Resources Center of the University of Illinois at Chicago. Schematics were created with BioRender.com.

## AUTHOR CONTRIBUTIONS

JR conceptualized the project. YMK and JR developed the overall study design and supervised the study. YMK, GM, SC, and PG(Gajwani) performed the experiments and interpreted the experimental data, XW and MAS performed transcriptomic data analysis, PTT, SV and SSYE performed the image analysis, YMK and JR wrote the initial manuscript draft. GM and PG (Grippo) generated KPC mice. TVN and KVN evaluated human samples. All authors provided feedback and revisions for the manuscript.

