## Extended Data for "Impaired Barrier Integrity of the Skeletal Muscle Vascular Endothelium Drives Progression of Cancer Cachexia"

### Extended Data Figures and Legends

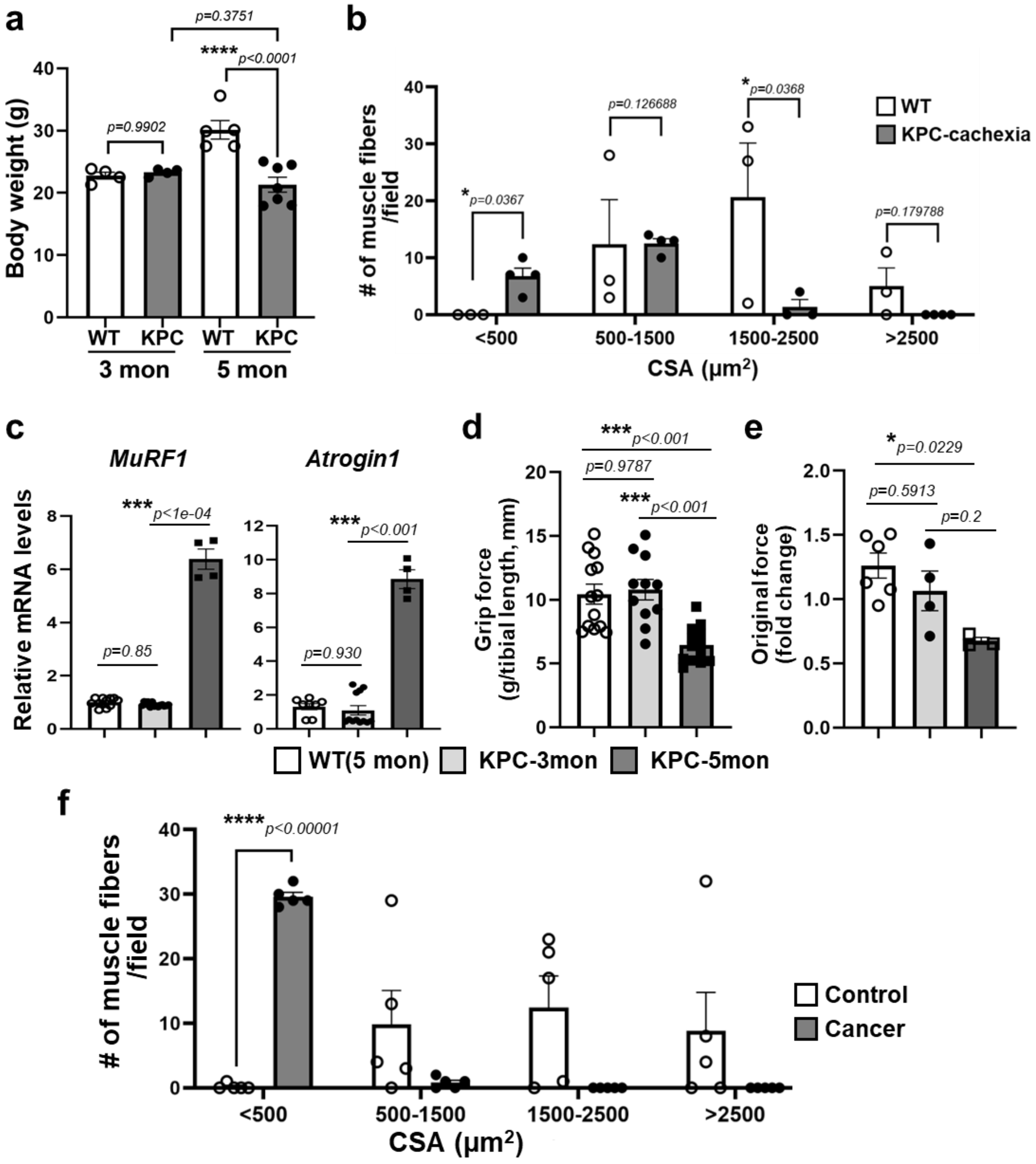


**Extended Data Fig. 1. Cachexia phenotype in KPC-cachexia mice and in cancer patients.** **a.** Body weight of control (WT, 3 and 5 month) and KPC mice (3 and 5 month). We used age matched C57/BL6J mice as control (WT). The statistical analysis was performed by one-way ANOVA, followed by post-hoc multiple comparison analysis using Tukey test, and the adjusted p-values were reported. **b.** The number of muscle fibers in WT and KPC-cachexia mice was sorted by CSA value. The area of muscle fibers was distributed mainly 1500-2000 $\mu m^2$. Thus, we considered that below 500 $\mu m^2$ fibers are small and over 2500 $\mu m^2$ fibers are big. Data are mean±SE for n=5-7 mice. The *p* values were evaluated by multiple unpaired t-test. **c-e.** Characteristics of cachexic syndrome in control (WT) and KPC mice (3 and 5 month). **c.** Expression of cachexia markers, *MuRF1* and *Atrogin1* in tibialis anterior (TA) muscles by RT-qPCR. Data are mean± SE for n=4-11 mice. The statistical analysis was performed by one-way ANOVA, followed by post-hoc multiple comparison analysis using Tukey test, and the adjusted p-values were reported. **d.** The four limbs grip strength of mice was measured average five times for each mouse using a grip strength test meter and were normalized by tibial length. Data are mean±SE for n=3-6 mice. The statistical analysis was performed by one-way ANOVA, followed by post-hoc multiple comparison analysis using Tukey test, and the adjusted p-values were reported. **e.** The fatigue of mice was evaluated using the values of grip force in **d** and presented with fold change of decrease of original force. The statistical analysis was performed by one-way ANOVA, followed by post-hoc multiple comparison analysis using Tukey test, and the adjusted p-values were reported. **f.** The number of human muscle fibers was sorted by CSA value. We considered that below 500 $\mu m^2$ fibers are small and over 2500 $\mu m^2$ fibers are big. Data are mean±SE for n=5-6 patients. The *p* values were evaluated by multiple unpaired t-test.

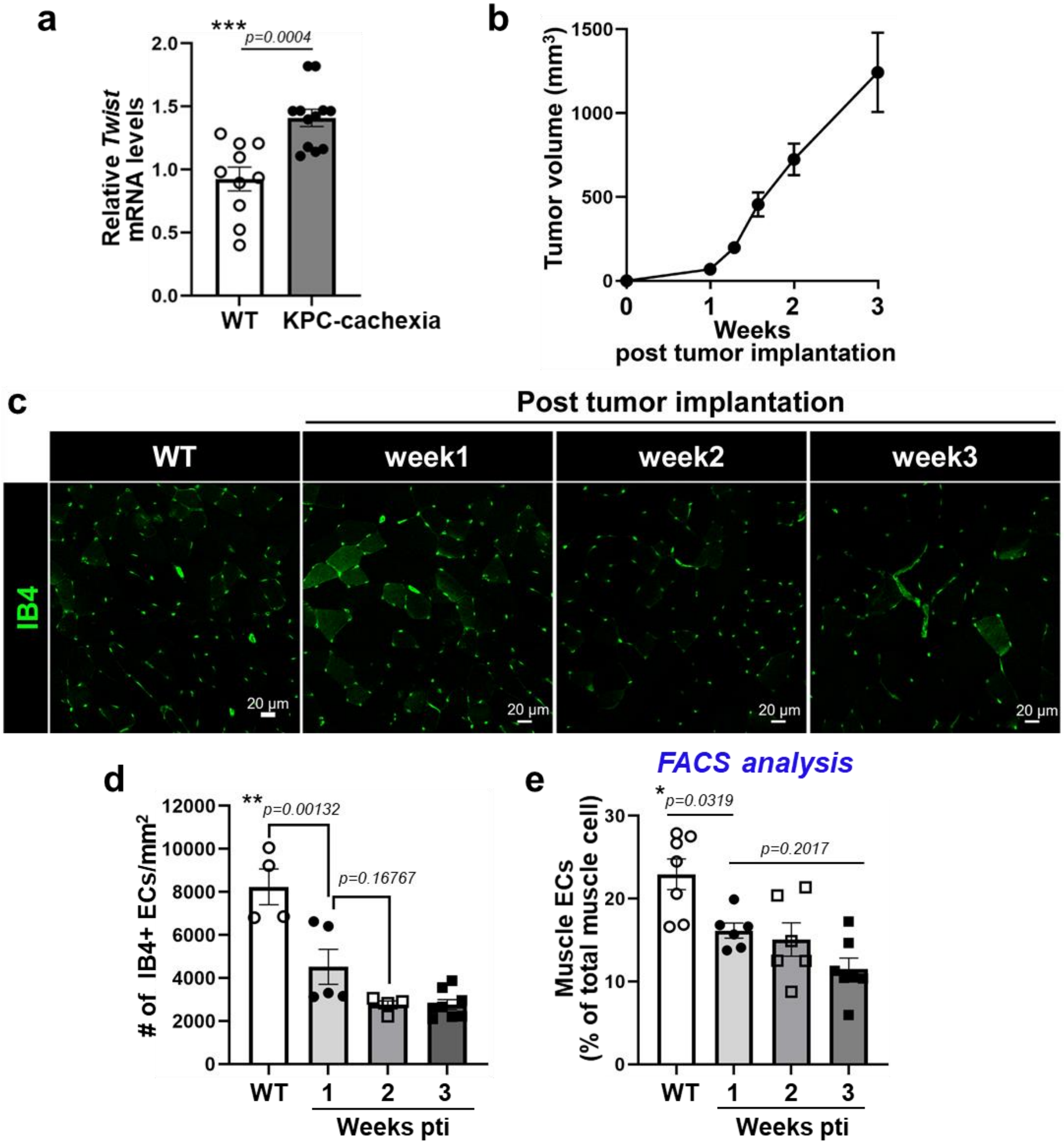


**Extended Data Fig. 2. Muscle vascular density decreases in a melanoma model of cachexia. a.** Mixed skeletal muscles (TA, GC, and quadriceps) from WT and KPC-cachexia mice were harvested and enzymatically digested and muscle ECs were isolated with CD31 microbeads and extracted total RNA. The expression of EndMT gene, *Twist* was determined by RT-qPCR with specific primers. The mRNA levels of *Twist* were normalized by *PPIA* levels and presented fold change. Data are mean±SE for n=10-12 biological replicates. The *p* values were evaluated by unpaired, two-tailed t-test. **b-e.** Mouse melanoma B16F10 cells ($10^6$ cells/100 µL PBS) were subcutaneously implanted in the dorsal flank of C57/BL6 mice and the mice were evaluated on 1, 2, and 3 weeks. Control mice were injected with PBS. **b.** Tumor growth post tumor implantation (pti). **c.** Cross-sectioned gastrocnemius muscles from WT mice and melanoma bearing mice were stained with IB4 for ECs (green). The images were taken using confocal microscopy (LSM880, 20x objective). **d.** Quantification of muscle vasculature density in **c** was presented with number of IB4+ ECs per muscle area ($mm^2$). Data are mean±SE from n=4-7 mice. The statistical analysis was performed by one-way ANOVA, followed by post-hoc multiple comparison analysis using Tukey test, and the adjusted p-values were reported. **e.** Mixed skeletal muscles (TA, GC, and quadriceps) from WT and melanoma bearing mice were harvested and enzymatically digested and the same number of total cells were stained with CD31 and CD45 antibodies and further stained with DAPI for FACS analysis. The CD31**+**CD45**-**DAPI**-** cells were considered as live pure ECs and presented with percent (%) of ECs in total muscle cells. Data are mean±SE for n=6-7 mice. The statistical analysis was performed by one-way ANOVA, followed by post-hoc multiple comparison analysis using Tukey test, and the adjusted p-values were reported.

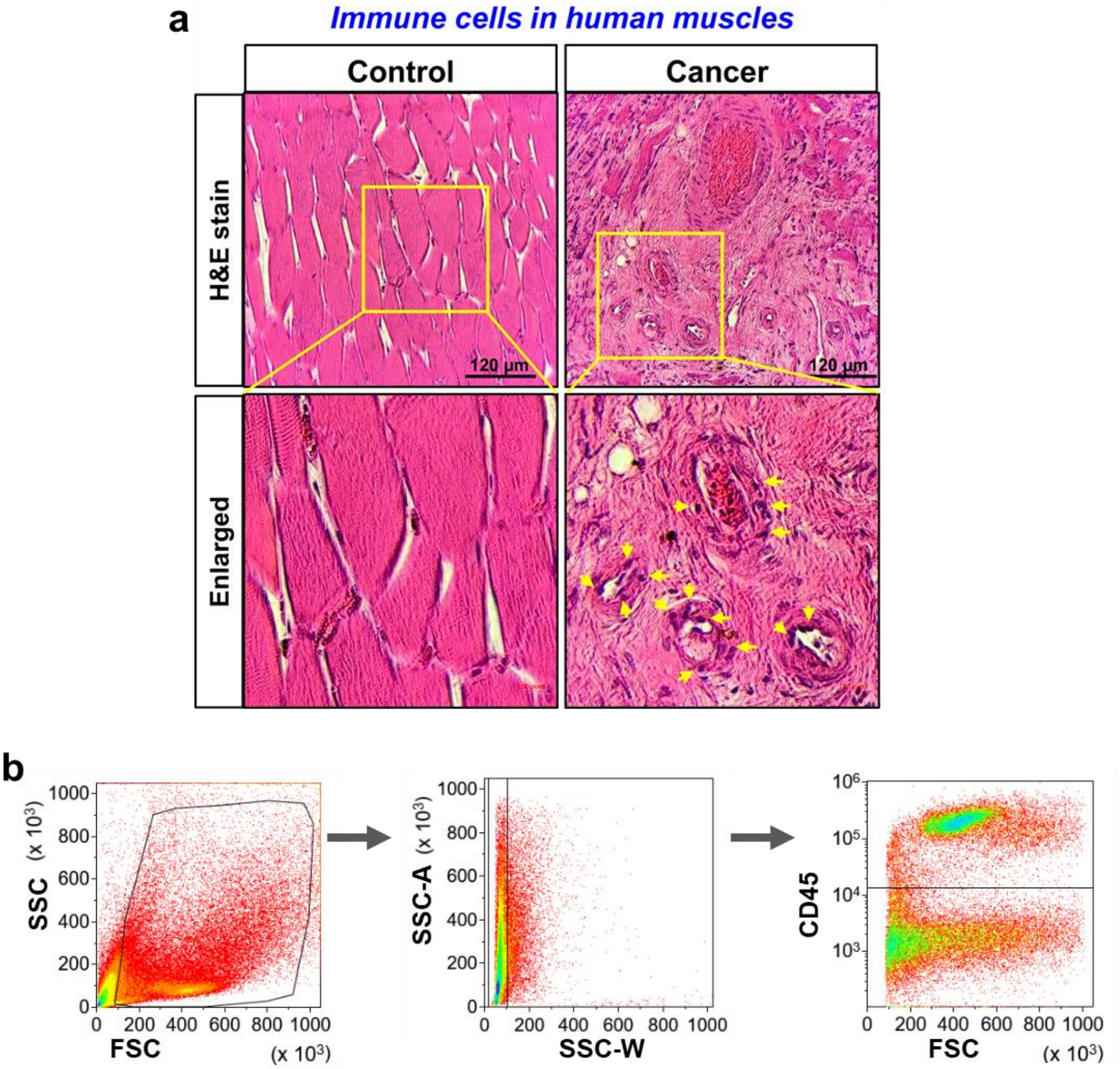


**Extended Data Fig. 3. Decrease of muscle vascular density is associated with the increase of infiltration of immune cells in cachexic muscles. a.** Abdominal muscles from control patients and cancer cachexia patients were stained with H&E and evaluated immune cell infiltration. The yellow arrow heads in enlarged box indicate the infiltrated immune cells in near blood vessels. **b.** Gating strategy for FACS analysis at **Main Figs. 3f-g**.

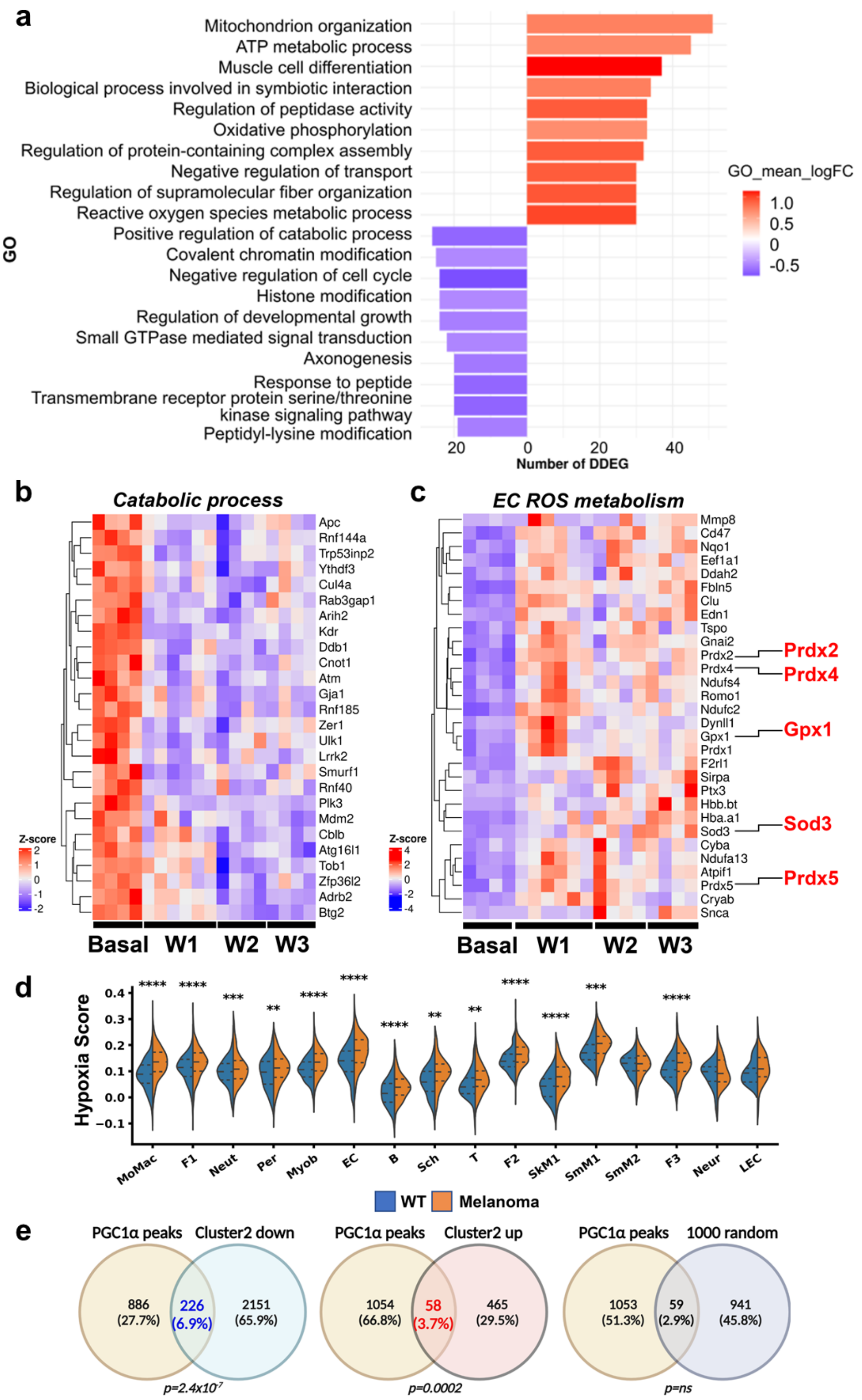
a
Mitochondrion organization
ATP metabolic process
Muscle cell differentiation
Biological process involved in symbiotic interaction
Regulation of peptidase activity
Oxidative phosphorylation
Regulation of protein-containing complex assembly
Negative regulation of transport
Regulation of supramolecular fiber organization
Reactive oxygen species metabolic process
Positive regulation of catabolic process
Covalent chromatin modification
Negative regulation of cell cycle
Histone modification
Regulation of developmental growth
Small GTPase mediated signal transduction
Axonogenesis
Response to peptide
Transmembrane receptor protein serine/threonine kinase signaling pathway
Peptidyl-lysine modification
GO
GO_mean_logFC
1.0
0.5
0
-0.5
20
0
20
40
Number of DDEG
b
Catabolic process
Apc
Rnf144a
Trp53inp2
Ythdf3
Cul4a
Rab3gap1
Arih2
Kdr
Ddb1
Cnot1
Atm
Gja1
Rnf185
Zer1
Ulk1
Lrrk2
Smurf1
Rnf40
Plk3
Mdm2
Cblb
Atg16l1
Tob1
Zfp36l2
Adrb2
Btg2
Z-score
Basal
W1
W2
W3
c
EC ROS metabolism
Mmp8
Cd47
Nqo1
Eef1a1
Ddah2
Fbln5
Clu
Edn1
Tspo
Gnai2
Prdx2
Prdx4
Ndufs4
Romo1
Ndufc2
Dynll1
Gpx1
Prdx1
F2rl1
Sirpa
Ptx3
Hbb.bt
Hba.a1
Sod3
Cyba
Ndufa13
Atpif1
Prdx5
Cryab
Snca
Prdx2
Prdx4
Gpx1
Sod3
Prdx5
Z-score
Basal
W1
W2
W3
d
Hypoxia Score
MoMac
F1
Neut
Per
Myob
EC
B
Sch
T
F2
SkM1
SmM1
SmM2
F3
Neur
LEC
WT
Melanoma
e
PGC1α peaks
Cluster2 down
886 (27.7%)
226 (6.9%)
2151 (65.9%)
p=2.4x10-7
PGC1α peaks
Cluster2 up
1054 (66.8%)
58 (3.7%)
465 (29.5%)
p=0.0002
PGC1α peaks
1000 random
1053 (51.3%)
59 (2.9%)
941 (45.8%)
p=ns

**Extended Data Fig. 4. Transcriptomic analysis of muscle cells (ECs and non-ECs). a-c.** Unbiased bulk RNA-seq analysis of dynamic differentially expressed genes in muscle vascular ECs during distal tumor growth. **a.** Top 10 GO pathways in muscle ECs. List of genes for catabolic process (downregulated, **b**) and EC ROS metabolism (upregulated, **c**). **d-g.** Unbiased scRNA-seq analysis of differentially expressed genes in muscle non-ECs (mixed muscle cells) during distal tumor growth. **d.** Hypoxia score of total mixed muscle cells from WT and melanoma bearing mice. The *p* values were evaluated by pairwise Mann-Whitney U test for each cell type. ***p* < 0.01, ****p* < 0.001, ****p* < 0.0001. MoMac; mono-macrophage,F1; fibroblast1, F2; fibroblast1, F3; fibroblast3, Neut; neutrophil, Per; pericyte, Myob; myoblast, EC; endothelial cell, B; B cell, Sch; schwann cell, T; T cell, SkM1; skeletal muscle 1, SkM2; skeletal muscle 2, smM1/2; smooth muscle cell, Neur; neuronal nerve, LEC; lymphatic endothelial cell. **e.** Overlap between genes adjacent to PGC1α ChIPseq peaks (n= 1112) and differentially expressed genes in the unique tdTomato positive (+) EC cluster 2 (left and middle) as compared to the average overlap of 1000 random subsets of 1000 genes from the background genes (n= 15960) (right) (hypergeometric test).

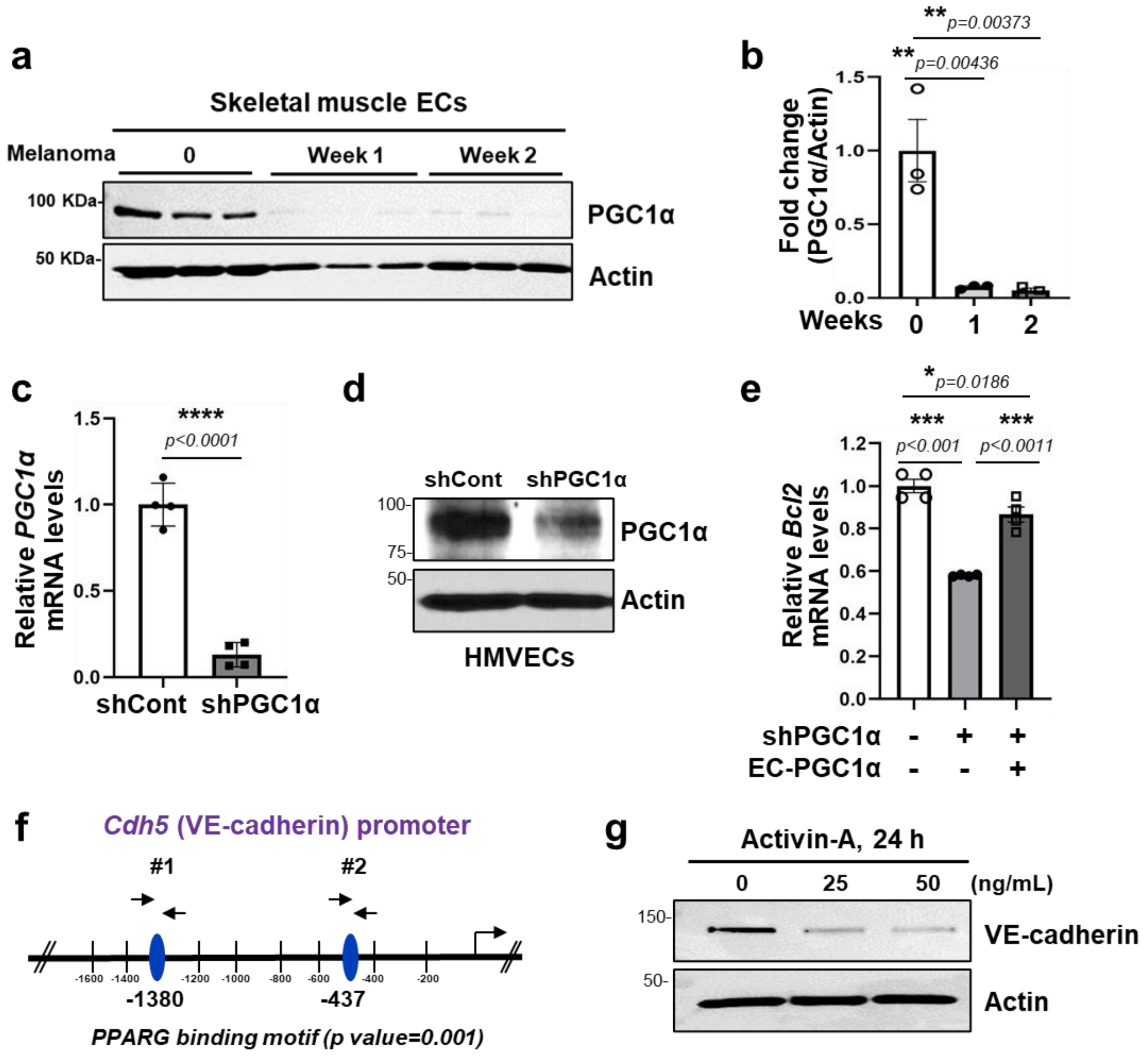


**Extended Data Fig. 5. Endothelial PGC1α is critical for endothelial cell viability. a**. The mixed muscles (GC, TA, and quadriceps) from control and melanoma bearing mice for 1 or 2 weeks were harvested, enzymatically digested, and isolated muscle ECs using CD31-conjugated microbeads. The PGC1α expression was determined by Western blotting with specific antibody. Each lane represents one mouse. **b**. The levels of PGC1α in **a** were normalized by actin and presented fold change. Data are mean±SE for n=3 mice. The statistical analysis was performed by one-way ANOVA, followed by post-hoc multiple comparison analysis using Tukey test, and the adjusted p-values were reported. The original blots can be found in Source data. **c-d.** HMVECs were treated with lenti-viral shRNA for Control and PGC1α for 72 h. **c.** The mRNA levels of PGC1α by RT-qPCR. Data are mean±SE for n=4 biological replicates. The *p* values were evaluated by unpaired, two-tailed t-test. **d.** The protein levels of PGC1α by Western blotting. The representative blots are from 3 independent experiments. The original blots can be found in Source data. **e.** HMVECs were co-transfected with lenti-viral shRNA for PGC1α and PGC1α plasmid for 72h. The mRNA levels of *PGC1α* and *Bcl2* were determined by RT-qPCR, normalized by *18s* levels, and presented with fold change. Data are mean±SE for n=4 biological replicates. The statistical analysis was performed by one-way ANOVA, followed by post-hoc multiple comparison analysis using Tukey test, and the adjusted p-values were reported. **f.** The putative PPARγ binding motifs in *Cdh5 (VE-cadherin)* promoter using EPD (eukaryotic promoter database, https://epd.epfl.ch//index.php). The #1 (-1400 bp ~ -1300bp) and #2 (-500 bp ~ -400bp ) motifs were evaluated by ChIP-qPCR. **g.** The serum starved ECs were treated with Activin A (25 and 50 ng/mL) for 24 h. The cell lysate was determined for VE-cadherin expression by Western blotting with VE-cadherin specific antibody. The representative blots are from 3 independent experiments.

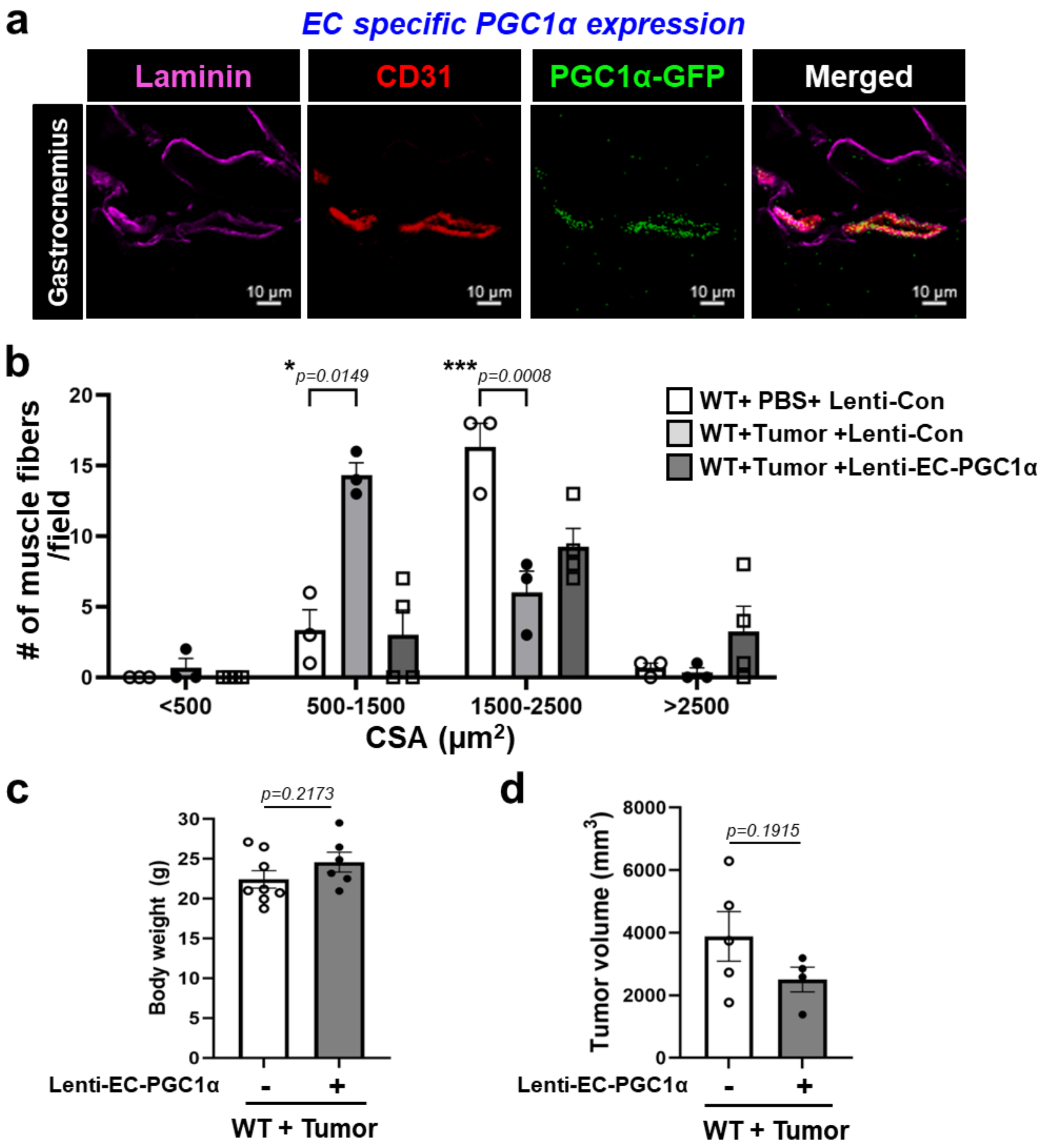


**Extended Data Fig. 6. Endothelial specific PGC1α overexpression in muscle vasculature prevents muscle dysfunction of tumor bearing mice. a-c.** Melanomas were subcutaneously implanted in the dorsal flank of WT mice and then the mice were intramuscularly received with lenti-control and lenti-EC-PGC1α-GFP virus. Control mice were received with same volume of PBS. All mice were examined in 3 weeks after tumor implantation. **a**. The cryosections of gastrocnemius were stained with EC-specific CD31 antibody (red) and muscle base membrane laminin antibody (magenta). Exogenously overexpressed EC-PGC1α was co-expressed with GFP. The images were taken using confocal microscopy (LSM880, 20x objective) and the colocalization of CD31 (red) and GFP (green) but not with laminin indicates specific expression of EC-PGC1α in muscle vasculature. **b.** The number of muscle fibers was sorted by CSA value. We considered that below 500 $\mu m^2$ fibers are small and over 2500 $\mu m^2$ fibers are big. Data are mean±SE for n=3-4 mice. The *p* values were evaluated by Tukey multiple comparison t-test. **c.** The effect of intramuscularly overexpressed EC-PGC1α on body weight of tumor bearing mice. Data are mean±SE from n=6-8 mice. The *p* values were evaluated by unpaired, two-tailed t-test. **d.** The effect of intramuscularly overexpressed EC-PGC1α on tumor growth. Data are mean±SE from n=4-5 mice. The *p* values were evaluated by unpaired, two-tailed t-test.
